# Absence of cortical reorganization following an internal capsule stroke in rodents

**DOI:** 10.1101/2025.10.31.685962

**Authors:** Patrick E. Cettina, David J. Guggenmos, Siddharth S. Sivakumar, Maxwell D. Murphy, H. Scott Barbay, Randolph J. Nudo, David T. Bundy

## Abstract

Stroke is a leading cause of disability, causing chronic motor impairments in many survivors. Although recovery is correlated with cortical reorganization, the impact of lesion location on post-stroke reorganization is uncertain. We compared movement-related neural activity following experimental infarcts to the motor cortex (M1) or internal capsule in rats. Neural activity was recorded from motor and somatosensory regions during a skilled pellet retrieval task longitudinally during the course of recovery. Cortical lesions resulted in early behavioral recovery accompanied by widespread reductions in neural activity across ipsilesional regions, indicative of large-scale reorganization. In contrast, internal capsule lesions produced delayed recovery with no evidence of long-term ipsilesional cortical reorganization. These findings challenge the relevance of cortical reorganization for recovery from subcortical lesions and indicate that post-stroke recovery mechanisms are lesion-specific and that models targeting subcortical white matter are essential for maximizing translational relevance.

## 1. Introduction

Stroke is the most common neurological disorder in the United States with nearly 800,000 new cases annually and a total prevalence of ∼9.4 million American stroke survivors (Martin et al., 2025). Although most patients survive their initial infarct, stroke is the leading cause of long-term disability due to cognitive and functional deficits that occur in the acute and chronic phases (Jackson & Chari, 2019). In particular, ∼40% of stroke survivors experience chronic motor deficits impacting their ability to perform activities of daily living (Cramer et al., 1997; E. Mayo et al., 1999). Although rehabilitation can drive increases in motor function, those gains typically plateau at 3-6 months post-stroke (E. Mayo et al., 1999; Lang et al., 2021). Therefore, there is a significant need to develop new therapies to restore function after stroke.

Importantly, a large body of work has examined the relationship between neuroplasticity and motor recovery. Preclinical models of stroke have been utilized to show alterations in motor and sensory maps in both perilesional and distant sensorimotor regions (Frost et al., 2003; Kraft et al., 2018; Nishibe et al., 2015; Nudo et al., 1996; Nudo & Milliken, 1996) . Similarly, interhemispheric functional connectivity is decreased initially, with restoration associated with motor recovery (Bauer et al., 2014; Hakon et al., 2018; Quattromani et al., 2018; Van Meer et al., 2010). However, while rehabilitation can increase the extent of remapping, the time course of motor recovery is distinct from the time course of cortical remapping, calling into question the specific role of cortical remapping in recovered movements (Eisner-Janowicz et al., 2008; Nishibe et al., 2015; Plautz et al., 2023). Along with preclinical studies, observations in clinical patients have also found that post-stroke recovery is associated with alterations in the location and extent of task-related neural activity (Cramer et al., 1997; Ward, 2003; Ward et al., 2003; Willer et al., 1993), stimulation-induced motor maps (Fridman, 2004), and interhemispheric functional connectivity (Carter et al., 2010, 2012). However, results are often heterogeneous across patients. Therefore, we still lack an understanding of the causal relationship between neuroplasticity and post-stroke recovery.

This gap in knowledge is compounded by the variability in stroke etiology, size, and location. Although most strokes (∼85%) are ischemic (Murphy & Werring, 2020), the location and extent of infarcts are variable across patients, extending into either or both cortical and subcortical structures in different patients (Corbetta et al., 2015; Favaretto et al., 2022; Kang et al., 2003; Stolz et al., 2006). Despite this variability, damage to the internal capsule is of particular importance due to the association between white matter damage within the corticospinal tract (CST) and post-stroke motor impairment (Carter et al., 2010; Corbetta et al., 2015; Marin & Carmichael, 2018; Y. Wang et al., 2016; Yamashita et al., 2016). Furthermore, when only patients with subcortical strokes are examined, limited alterations in functional connectivity are observed, calling into question the role and necessity of cortical reorganization in motor recovery and highlighting the need for a more mechanistic understanding of the patient-specific role of cortical reorganization in stroke recovery (Branscheidt et al., 2022; Makin & Krakauer, 2023).

Despite the relevance of white matter integrity to functional deficits, common preclinical models used for examining the mechanisms of post-stroke plasticity do not typically model the location and extent of strokes that typically occur in patients (Edwardson et al., 2017; Marin & Carmichael, 2018; Sozmen et al., 2012). Therefore, we still lack a mechanistic and location-specific understanding of the role of neuroplasticity in post-stroke recovery. Importantly, recent work has sought to develop novel preclinical models of subcortical stroke targeting the internal capsule (Blasi et al., 2015; Frost et al., 2006; Kim et al., 2014, 2021; Murata & Higo, 2016; Song et al., 2017; Wen et al., 2019a), enabling studies of the lesion-specific role of neuroplasticity after stroke. Although early studies have shown motor deficits associated with subcortical models of stroke, the specific changes in cortical activity associated with subcortical stroke remain uncertain.

In this study we sought to address this gap in knowledge by comparing the alterations movement-related neural activity following recovery from rodent models of cortical and white matter stroke. Following training to perform an automated skilled single-pellet retrieval task (Bundy et al., 2019), rats were randomized to receive a focal ischemic infarct to the forelimb representation within the primary motor cortex (caudal forelimb area; CFA) or a stimulation-guided infarct to the internal capsule followed by implantation of chronic microelectrode arrays to M1 (CFA), the rostral forelimb area (RFA; a motor region similar to premotor cortex in non-human primate species) (Sievert & Neafsey, 1986; Rouiller, 2009), and S1. We then recorded neural firing during reaching task performance throughout the course of spontaneous recovery from infarcts. Our initial hypothesis was that each lesion would be associated with distinct patterns of post-stroke reorganization. Following an infarct to M1 (CFA), we observed alterations in task-related neural activity in surrounding regions. Following a capsular infarct, however, an absence of ipsilesional cortical reorganization was observed. Together these findings support that recovery from lesions in distinct locations may be facilitated by distinct neural mechanisms.

## 2. Methods

### 2.1 Experimental Design

All procedures were approved by the University of Kansas Medical Center Institutional Animal Care and Use Committee. All experiments were performed in young adult (4-6 months old) Long-Evans rats (*Rattus norvegicus*). Fourteen rats were randomized into either a control group or one of two experimental stroke groups that received a focal ischemic infarct to M1 (CFA) or a focal ischemic infarct targeting the internal capsule (IC). To evaluate the changes in movement-related neural activity associated with post-stroke recovery, chronic neural recordings were obtained from motor cortex (M1, CFA), premotor cortex (RFA), and primary somatosensory cortex (S1) during performance of a skilled reaching task for eight weeks following the experimental infarct.

### 2.2 Skilled Pellet Retrieval Task

Rats were trained to perform a skilled pellet retrieval task within a custom-designed automated behavior box that we have described previously (Figure 1) (Bundy et al., 2019). Rats were placed within a 12” x 12” (305 mm x 305 mm) acrylic box with 15 mm slits on each side of the front and back walls limiting forepaw usage to the preferred forepaw. In each trial, rats were initially required to press a button at the back of the box, causing a door diagonally opposite at the front of the box to open providing access to a food reward on a shelf placed outside of the box. Rats then transitioned to the front of the box and performed a reach-to-grasp movement to retrieve a 45 mg banana pellet (Bio-serv) from a shelf outside of the box. An infrared beam break was placed across the front of the box to detect reaching attempts. The door was closed following completion of a reach attempt, providing a well isolated reaching trial for alignment to neural data. A trial was aborted if the rodent did not attempt to retrieve the pellet within 20 seconds of the door opening. The pellet retrieval portion of the task has been well validated for assessment of forepaw function in rodents in experimental stroke and TBI models (Corbett et al., 2017; Nishibe et al., 2010, 2015; Whishaw & Pellis, 1990).

**Figure 1.**
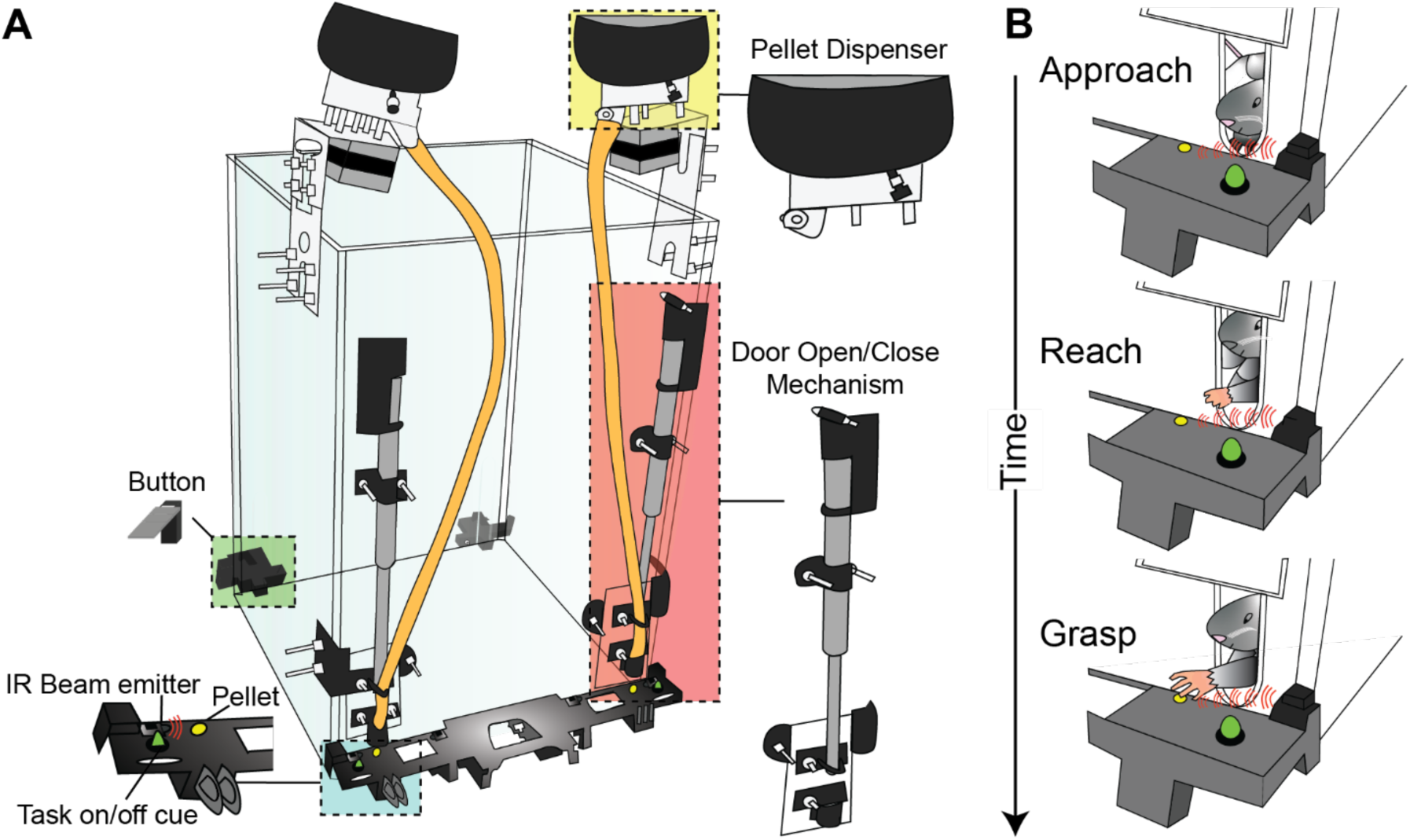
Automated single-pellet retrieval task. **A.** Behavioral recovery and task-related neural activity were assessed using a single-pellet retrieval task. A custom-made automated behavior box controlled the task flow. Each trial is initialized with a button press by the preferred forepaw at the back of the box, which triggers a door at the front to open providing access to a food reward placed on a shelf placed outside of the box. Rats are then required to cross diagonally the front of the box to perform a reach-to-grasp movement with the preferred forepaw to retrieve the food reward. An IR beam break detects the completion of the initial retrieval attempt and triggers the door to close to limit repeated attempts. An LED light was used to synchronize neural data with video recordings to isolate behavioral events. **B.** Video recordings of task performance were used to manually score each trial. Each reach-to-grasp trial was segmented into an ‘Approach’ or preparatory epoch prior to reach onset, a ‘Reach’ epoch beginning with the onset of the initial reaching movement, and a ‘Grasp’ phase initiating with the onset of grasp and including the retrieval movement.

Prior to training, we determined each rat’s preferred forelimb. All training and post-stroke behavioral assessments were performed with the identified forelimb. Rodents were trained under a progressive-associative training paradigm. Initially, rats were trained to perform reaching movements and were habituated to the door opening and closing between attempts. Next, to train rodents to associate the button press with the reward, pellets were dispensed to reward each successful button press. After initiating consistent button press attempts, rats were finally trained to perform the full task and receive their pellet reward by completing the reach-to-grasp task following successful button presses. Rats that achieved >50% successful trials for 3 consecutive sessions of 25 trials per session were included in the study. Task performance was maintained with 2-3 training sessions per week until surgery could be performed (typically 1-2 weeks).

### 2.3 Surgical Methods

#### General Procedures

All surgical procedures were performed under aseptic conditions. Rats were initially anesthetized with isoflurane (to effect) followed by ketamine (80-100mg/kg, i.p.) and xylazine (5-10 mg/kg, i.m.) and anesthesia was maintained with supplemental injections of ketamine (10-20 mg/hr i.m.). Rodents were placed in a stereotaxic frame, an incision was made along the midline of the scalp, and the temporalis muscle was resected. A laminectomy was performed to reduce cortical swelling during the craniectomy. A craniectomy was then made over the sensorimotor cortex contralateral to the preferred forelimb and the dura was retracted. The caudal forelimb area (CFA/M1) and rostral forelimb area (RFA/PM) were then mapped using intracortical microstimulation (ICMS) as described previously (Nishibe et al., 2010, 2015). Following surgery, rats were administered penicillin (45,000 IU s.c.) to limit infection risk. Four doses of buprenorphine (0.05-0.1 mg/kg subcutaneously) and acetaminophen (20-40 mg/kg orally) were administered over 48 hours as analgesics.

### Stroke Models

Rats were randomized using a block randomization scheme into healthy control (n=4), cortical infarct (n=5), and subcortical infarct (n=5) groups. All infarcts were made using a photothrombotic model of focal ischemia (Figure 2). After the photoactive dye rose Bengal is injected intravenously and activated using green light (520 nm), a thrombus is formed through platelet accumulation (Bergeron, 2003; Watson et al., 1985). Photothrombotic models of stroke have been used to produce consistent and focal ischemia in rats (Moon et al., 2009; Schrandt et al., 2015; Watson et al., 1985), mice (Lee et al., 2007), and non-human primates (Furuichi et al., 2003; Ikeda et al., 2013; Maeda, Moriguchi, et al., 2005; Maeda, Takamatsu, et al., 2005; Sawada et al., 2014; Tomizawa et al., 2015). For the cortical infarct group, lesions were targeted to CFA/M1 following ICMS mapping identification. A laser-diode light source emitting green light (520 nm, Doric Lenses) was focused on CFA using a stereotaxically-mounted fiber optic placed 1-2 mm above the surface of the cortex. Rose Bengal was then administered (20 mg/kg i.v.) through the lateral tail vein and cortex was illuminated for 15 minutes (Figure 2A). Subcortical infarcts were targeted to the internal capsule using micro-stimulation guidance (Wen et al., 2019b). The internal capsule was localized, and lesions were generated with an optoelectric probe (Doric Lenses) consisting of a 50 µm tungsten wire extending 500 µm beyond a 70 µm fiber optic. Stereotaxic coordinates (-1 to -2 mm AP, 2 to 4 mm ML relative to bregma) were used to target the insertion of the optoelectric probe to the posterior limb of the internal capsule based upon a standard rat brain atlas (Paxinos & Watson, 2013). This range contained prominent representation of the internal capsule while allowing adjustment of the insertion site to avoid prominent blood vessels on the cortical surface. The location and depth (typically 5-7 mm ventral to the brain surface) of the internal capsule was confirmed with ICMS. Specifically, when the probe tip is in the internal capsule, ICMS produces forelimb movements at low current amplitudes (10-20 µA). After locating the internal capsule, rose Bengal (20 mg/kg i.v.) was administered, the laser diode light source was connected to the optoelectric probe tip, and light was delivered for 15 minutes to create a single subcortical infarct (Figure 2B).

**Figure 2.**
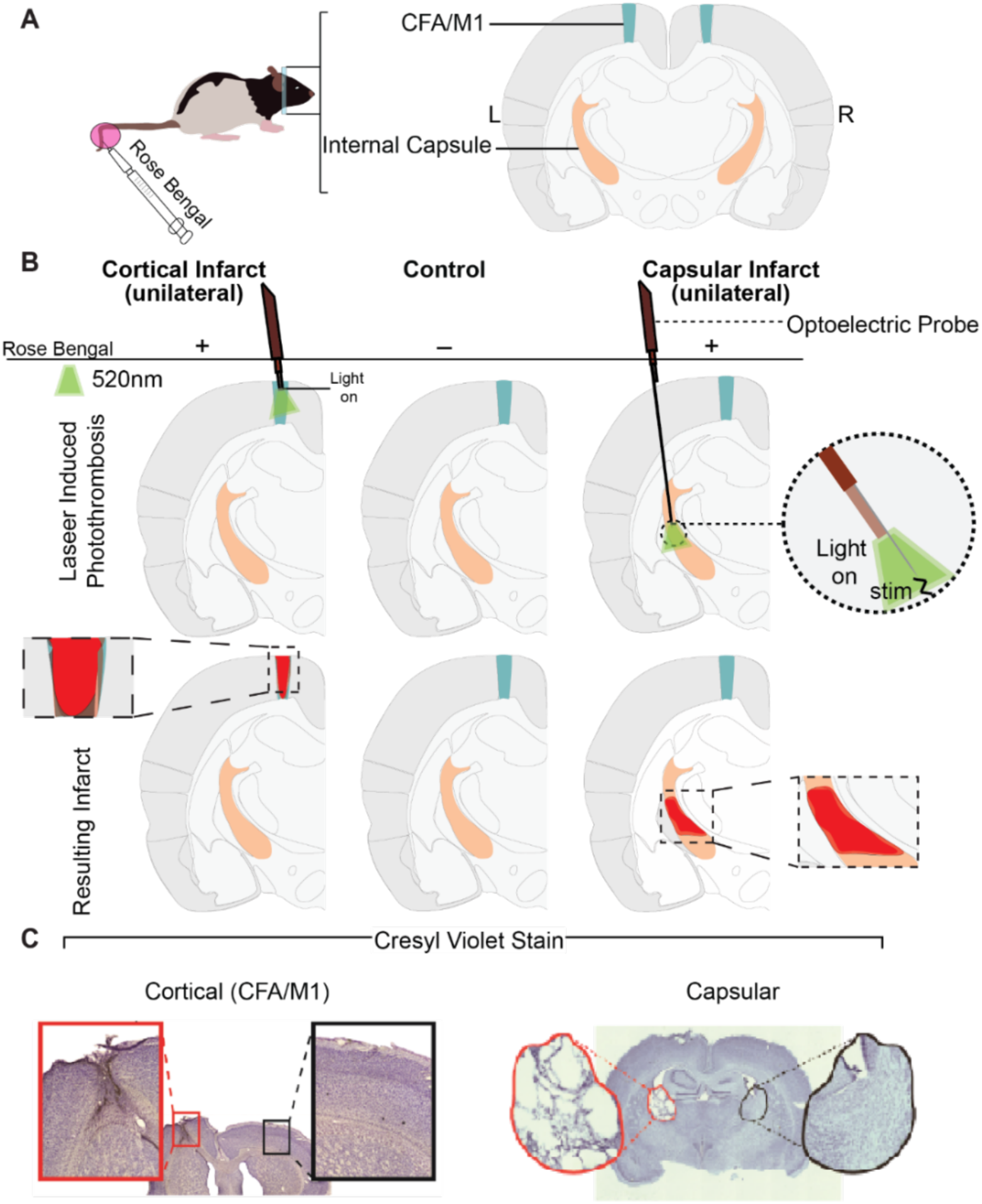
Specific targeting of photothrombotic infarcts. **A.** A photothrombotic technique was used to generate ischemic infarcts. A photoactive dye rose Bengal was administered with an IV tail vein injection and a focally targeted green light source was used to target ischemic infarcts to either the forelimb region of the primary motor cortex (CFA/M1) or the internal capsule (IC). **B.** Each rodent was randomized into one of three groups: Control, Cortical, Capsular. The hemisphere contralateral to the preferred forelimb of each rodent was targeted for both lesion groups. CFA and IC were each identified using stimulation mapping. For capsular infarcts both stimulation mapping and lesion induction were performed using an optoelectric probe ensuring accurate lesion targeting. Following injection of rose Bengal, the experimental groups were exposed to green (520nm) light at the target area (CFA or IC) for 15 minutes to induce focal ischemia. **C.** Exemplar lesions are shown in histological samples from rats not implanted with electrodes to limit tissue damage from removal of electrode arrays. Brains stained with cressyl violet for Nissl bodies show that infarcts were consistently and accurately targeted. Cortical lesions extended through all layers of cortex without impacting deeper subcortical structures (left). Capsular infarcts were accurately targeted to the IC with some limited infiltration of neighboring subcortical structures.

### Electrode Implants

Following infarcts, three 16-channel NiCr microwire electrode arrays (Microprobes) were implanted into RFA, CFA, and S1-forelimb contralateral to the preferred forepaw (Figure 3A). Probes were targeted to CFA and RFA using the ICMS maps described previously. Additionally, probes were targeted to the forelimb region of S1 using stereotaxic coordinates (-1.25 mm AP, +4.25 mm ML). All arrays were organized in 4x4 grids with 250 µm spacing between electrodes and included a reference electrode that was 500 µm shorter than the electrode contacts, allowing an impedance-matched reference in superficial layers of the cortex. A silver wire from each probe was attached to an 00-80 stainless steel skull screw to act as a ground. Electrode arrays were implanted to depths of approximately 1500 µm (Layer V) in RFA and CFA, and 1300 µm (layer IV) in S1 using a motorized micropositioner (Narishige International USA, Inc., Amityville, NY). Following electrode insertion, the cortex was covered with a silicone elastomer (Kwik-Cast, World Precision Instruments, Sarasota, FL), a head cap was constructed from dental acrylic *in situ*, and the scalp was sutured around the head cap.

**Figure 3.**
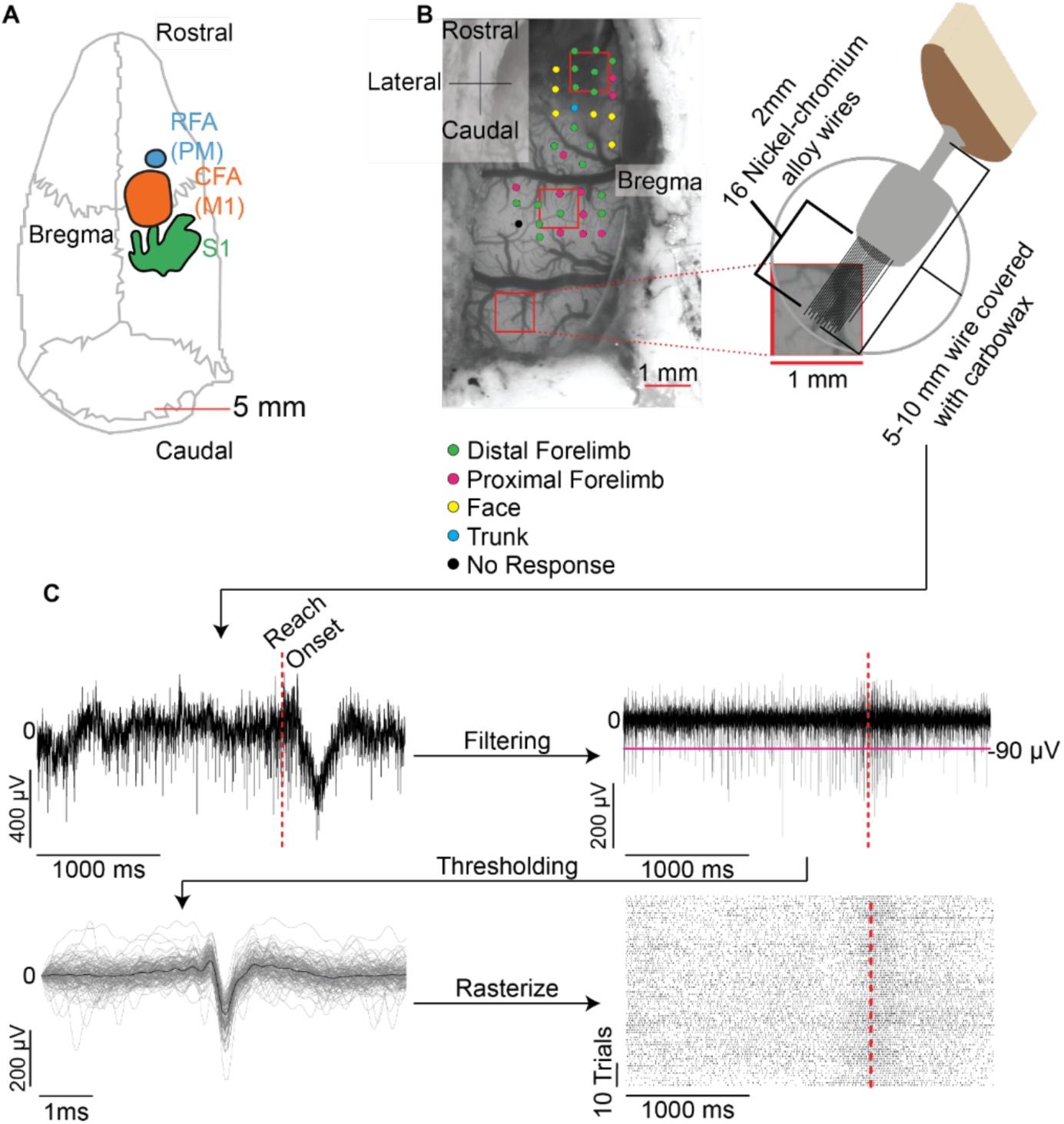
Electrode implants and neural data processing. **A.** Rats possess distinct primary (M1, caudal forelimb area, CFA) and secondary (premotor, rostral forelimb area, RFA) motor regions. Electrode implants were targeted into RFA, CFA, and the forelimb region of primary somatosensory cortex (S1). **B.** In each rat, three 16-channel Nickel-chromium microwire electrode arrays were implanted into RFA, CFA, and S1. Electrode implants were targeted to CFA and RFA using ICMS mapping. S1 electrode array implants were targeted using stereotaxic coordinates. **C.** Neural data from each recording session was filtered (300-3000 Hz), action potentials were detected with a threshold detector, and spikes were aligned across trials. Pre-processing and spike detection steps are shown for an exemplar recording from a single electrode.

### 2.4 Neurophysiological Data Collection

Task-related neural recording began 5 days post-surgery for each rat with two sessions of data collected each week for the following 8 weeks. On recording days, rodents were anesthetized using isoflurane to facilitate attachment of the recording headstages to each electrode array. Headstages (Intan RHD 2132, Intan Technologies, Los Angeles, CA) performed amplification and digitization at the rat’s head and were connected through a slip-ring commutator to an Intan biosignal amplifier (RHD2000, Intan

Technologies, Los Angeles, CA). Neural data was recorded at 20 kHz while rats were freely moving within the automated behavior box. Recordings ended after rats performed 50 trials or one hour had elapsed with a minimum recording duration of 30 minutes per day. Occasionally recordings were stopped and restarted or truncated due to head stages coming unplugged mid-session. One control rodent was excluded from neural data analysis because the headcap failed following only four sessions.

### 2.5 Behavioral Scoring

Behavioral information including button press times and infrared beam break readings were recorded as digital inputs by the Intan system to synchronize behavioral events with neural data. Additionally, video recordings of behavioral task performance were made from the front and back of the box with an LED used to synchronize neural data with videos. Behavioral videos were then manually scored to identify specific movement components within reach-to-grasp trials (Figure 1B). Because the single-pellet retrieval task is a well-standardized task to examine recovery in rodent models of stroke, neural and behavioral analyses focused on this task component. Specifically, reach onset was identified as the first video frame where the rodent lifted its forepaw to initiate a forward movement of the forepaw. Grasp onset was identified as the first frame where the rat’s distal forepaw began to contract. Although the door began to close immediately after completion of an initial reaching attempt, because a beam break was used to prevent the door from closing on their paw, rats occasionally made additional reaching attempts prior to the door closing. A trial was coded as successful if the rodent was able to retrieve the pellet into the box and a failure if they were unable to retrieve the pellet prior to the door closing or knocked the pellet off of the shelf. Successful trials were segregated into initial and delayed success trials depending upon whether the rodent retrieved the pellet with a single or multiple reach-to-grasp attempts. Trials without a reach attempt within the 20-seconds were considered aborted trials. Aborted trials and any trials attempted with the forepaw ipsilateral to the electrodes were excluded from analysis. Analyses of normalized success, normalized reach attempts, normalized reach-to-grasp interval, and trial counts were done using a one-way ANOVA and Tukey test for significance of pairwise group comparisons. To examine the trajectory of recovery and its relationship to each group a linear mixed effect model was built:

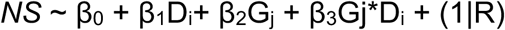

The LMM has a fixed and mixed effect structure where the main effects examined were for day (D_i_) and group (Gj: control, cortical, capsular) and a mixed effect of day*group was used to examine differences in the trajectory of behavioral recovery between groups with a random effect included for rodent-specific differences.

### 2.6 Neural Data Processing

#### Data Preprocessing

During daily recordings noisy channels were identified and noted for removal from later analysis. Initially data were rereferenced to the common average across all channels to remove common noise sources. Next, in order to examine neural firing activity, raw data was bandpass filtered between 300Hz-3000Hz using an elliptic IIR filter with the parameters: FS = 20,000; FSTOP1 = 250; FPASS1 = 300; FPASS2 = 3000; FSTOP2 = 3050; ASTOP1 = 70; APASS = 0.001; ASTOP2 = 70; MATCH = both. Processed data was then used to detect multi-unit action potentials.

#### Multi-unit Spike Detection

Following preprocessing, multi-unit spike events were detected. Spikes were detected using a threshold of -3.5x RMS voltage calculated for each electrode and recording session (Trautmann et al., 2019). Waveforms underwent visual inspection to remove noise, but no sorting into individual units was performed as multi-unit firing has been reported to be effective for neural dynamics analyses (Trautmann et al., 2019). Figure 3C shows the data processing steps for a single channel from an exemplar rat. Following multi-unit detection, firing rates were estimated for each channel through a Gaussian convolution of the binned raster data (spike instances binned into 1ms bins for the length of the time series).

### 2.7 Task-Related Neural Firing Analysis

Task-related neural data was initially processed as binary rasters aligned to the initiation of reaches. Action potentials were placed into 1 ms bins from 2000 ms before to 1000 ms after reach onset. Although data were not aligned to the grasp, all grasp attempts were contained within the 1000 ms post-reach period. As described above, a Gaussian convolution (σ=25 ms, kernel size=125 ms) was applied to the binary time series to estimate firing rate, and the data was stored for each trial. Baseline periods were collected using a +/- 1500 ms window around the temporal midpoint between trials to sample data from non-task periods. The same Gaussian convolution was applied to binned raster data during the baseline periods. The mean and standard deviation of baseline periods were used to normalize the firing rates during trial periods. Because the number of trials immediately after the lesion was limited in some rats, we specifically focused on neural activity following the plateau of behavioral recovery (weeks 5-8) to examine differences in the neural correlates of movement following recovery. Recording sessions with additional noise in the signals due to head stage displacement were removed from the analysis, leaving a minimum of six (75%) post-plateau recording sessions in each rat and resulted in approximately 262, 272, and 255 total trials on average for each rodent in the control, cortical, and capsular groups respectively. To balance the number of sessions considered between groups, we analyzed the six behavioral sessions within the post-plateau weeks 5-8 with the greatest number of trials for each rat.

### 2.8 Modeling Neural Firing Peri-Task

Finally, to estimate the separability of each condition/group, epoch, and region, a linear mixed model was used to relate these variables with neural firing in each rodent. The model was fit using the multi-unit spiking data described previously. To ensure consistent task-related neural activity and prevent differences in neural patterns associated with reduced success rate following the experimental infarcts, we limited analyses to trials where the rodent was successful. The data was then stored in a hierarchical format: Rodent → Session (day) → Region → Channels → Trials → Binned Neural Activity. The binned neural activity consisted of multi-unit spike counts within 50 ms bins from 2000 ms prior to reach onset to 1000ms after reach onset. The summed spike counts were then divided by 50ms to get the firing rate within each bin. Each temporal bin was then classified by task timing into an approach, reach-to-grasp execution, or retrieval period. Each epoch was encoded as a binary mask where 0 represented that time bin was not within the given epoch and a 1 represented when the time bin was within the given epoch. The reach-to-grasp execution was defined as the period between the initiation of reach and initiation of grasp in each trial. Importantly, only trials classified as successful following a single reach and grasp attempt were used in this model. The approach epoch was defined as the 2000 ms period before the reach-to-grasp execution. The retrieval phase was defined as the period from the onset of grasp until the end of the trial window 1000 ms after reach onset. All other variables were encoded as categorical variables including group (cortical lesion, capsular lesion, or control), region (RFA or S1), rodent ID, channel ID, and session ID. CFA was not included in the model to avoid biasing model fit because CFA was the targeted location of the infarct in the cortical group. The full equation for the linear mixed model is shown in equation 2:

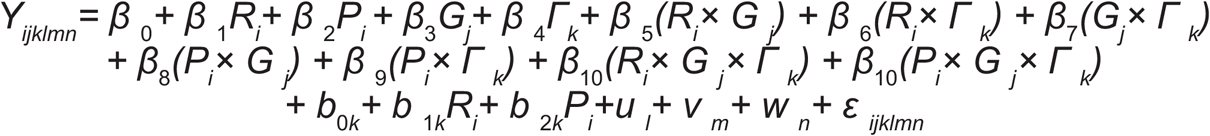

The LMM had a fixed effect structure where the main effects examined were for group (G_j_: control, cortical, capsular), epoch (R_i_, P_i_: reach, grasp), and region (Γ_k_: RFA, S1). The random effects specified in the model were for ChannelID in reference to both epoch, rodent, session ID, and trial ID. The purpose of including the random effects was to capture the excess hierarchical dependencies and heterogeneity of neural firing. The channel ID random effect specifically captured any variability in firing between individual channels, while the other random effects accounted for any variability inherent across animals, sessions, or individual trials. The model was fit using *MATLAB’s fitlme* function (Statistics and Machine Learning Toolbox). Following the construction of the model the goodness-of-fit was tested by gathering the R-squared between the original response vector Y and a reconstructed Y’ vector created by multiplying fixed effects weighs with their respective values for each bin. Finally, to test the impact of each fixed effect on its contribution to the fit model we nullified individual beta contributions and reconstructed the response vector in the same way as above. The difference in the capacity for each reduced model to reconstruct the original neural data relative to the full model was extracted and evaluated for each fixed effect.

## 3. Results

### 3.1 Cortical and capsular infarcts lead to distinct behavioral recovery trajectories

Although the presence of electrode arrays in ipsilesional sensorimotor cortices limited the ability to perform a histological comparison of lesion volumes, histological samples from exemplar rats show that both lesion models produced infarcts that were accurately targeted to CFA or the internal capsule respectively (Figure 2C). Furthermore, intraoperative examination of cortical infarct locations and sizes confirmed accurate targeting of cortical lesions to CFA. Additionally, each group had distinct patterns of behavioral deficits. Rodents in the control group maintained their level of task performance following electrode implantation, while both the cortical and capsular infarct groups showed a decrease in pellet retrieval success rate (one-way ANOVA: p < 0.001, F = 28; pairwise Tukey test control – cortical: p < 0.001, control – capsular: p < 0.001, cortical – capsular: p = 0.245), which plateaued 4-weeks post-surgery at a success rate below their pre-infarct baseline (Figure 4A). Although both the cortical and capsular groups plateaued at similar normalized success rates, their recovery trajectories are distinct. Specifically, the capsular group has a delayed recovery trajectory reaching the plateau phase later than the cortical infarct group (linear mixed effects model: Normalized Success ∼ Day*Group + (1|Rodent); one-way ANOVA on effects; intercept p < 0.001, F = 59; group p < 0.001, F = 12; day p = 0.95, F = 0.0039; group*day p < 0.001, F = 12) (Figure 4A). Additionally, the lesion groups also differ in post-infarct reaching speed, measured as the time interval from reach onset to grasp onset. While both groups show an initial decrease in reach speed, the capsular group recovers reaching speed faster with an average reach time similar to their pre-lesion performance by day 17 post-lesion despite worse success rates (Figure 4B). In contrast, the cortical group perform reaching movements with slower speed up to 31-35 days post-lesion. Despite differences in task performance between the lesion groups, the total number of reach attempts per trial are similar (Figure 4C). Although control rats perform more trials than either the cortical infarct or capsular infarct, both lesion groups increase the number of reaching attempts by 4-weeks post-surgery. Therefore, differences in task performance and task-related neural activity are not impacted by motivational differences or the number of trials included (Figure 4D). Furthermore, the separation between infarct groups on task success rate and reaching speed shows that behavioral recovery follows lesion-specific trajectories.

**Figure 4.**
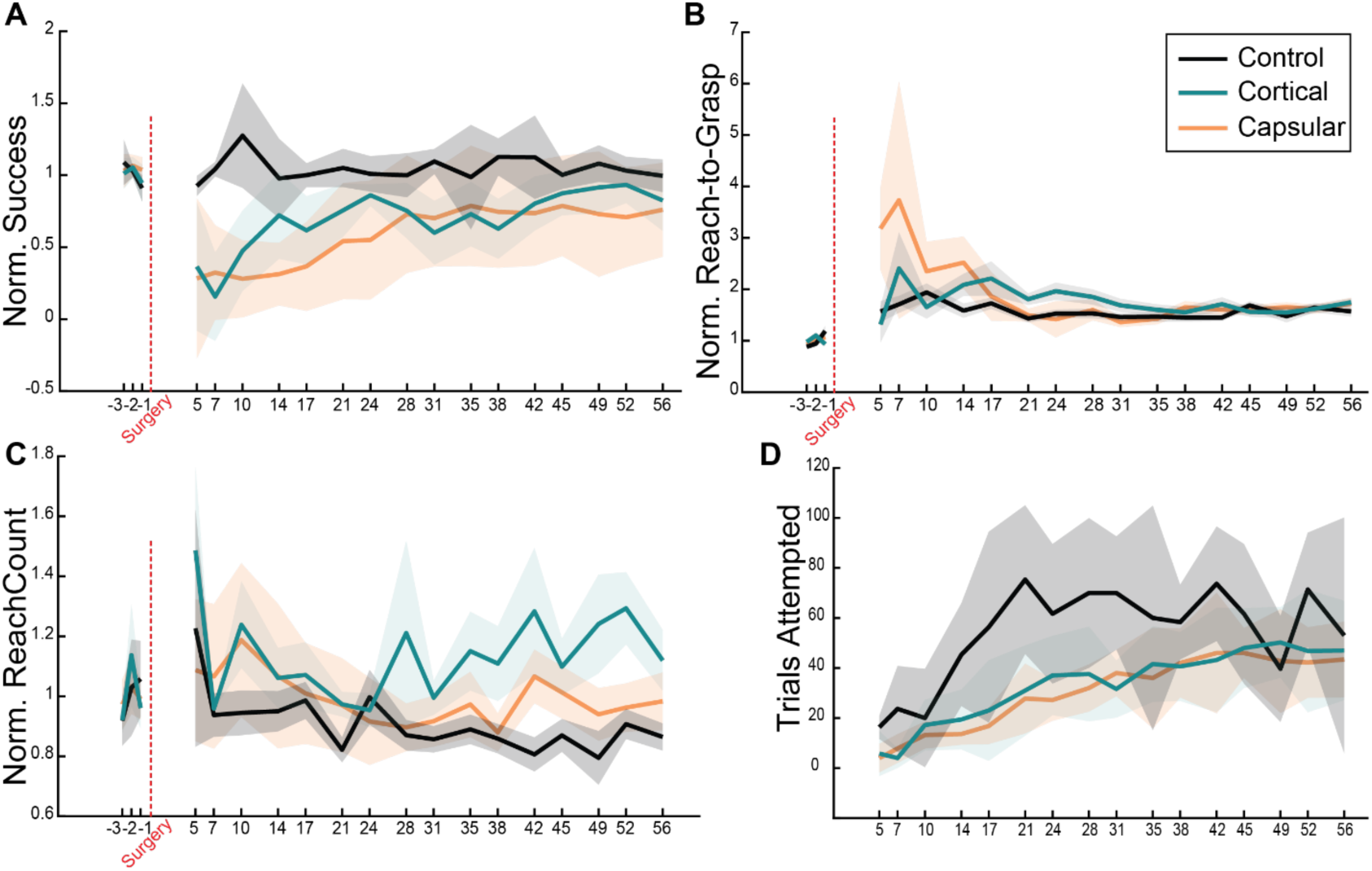
Lesion-specific behavioral recovery trajectories. **A.** Pellet retrieval success rates normalized to each rat’s (pre-surgical) baseline success rates. Control rodents significantly outperform the experimental groups (one-way ANOVA: p < 0.001, F = 28), and there is a separation in recovery trajectory dependent on stroke type (linear mixed effects model: Normalized Success ∼ Day*Group + (1|Rodent); one-way ANOVA on effects intercept p < 0.001, F = 59; group p < 0.001, F = 12; day p = 0.95, F = 0.0039; group*day p < 0.001, F = 12). **B.** Normalized reach-to-grasp interval encoded only on successful trials and gathered by acquiring the time of the last reach and grasp attempt within a trial if multiple occurred. Compared to healthy controls, both stroke groups perform reaching movements with slower speed (one-way ANOVA: p < 0.001, F = 21). Although all groups plateau to a similar reaching speed, the capsular group recovers movement speed about two weeks (days 17-28) earlier than the cortical group (one-way ANOVA: p < 0.001, F = 42). **C.** Normalized reach count per successful trial. Controls are successful using the fewest number of reaches (one-way ANOVA: p < 0.001, F = 70). **D.** Total number of trials attempted by each rodent group in each session. Each group takes ∼10 days to recover trial counts. Controls participated in a greater number of trials earlier following the surgery; however, the trial counts used for neural data analysis from the post-plateau period were not significantly different (one-way ANOVA: p < 0.001, F = 18; one-way ANOVA: p = 0.59, F = 0.52). All plots show mean +/- 95% confidence interval (CI).

### 3.2 Task-related neural activity is present across lesion groups

Following motor recovery task-related neural activity is observed from ipsilesional sensorimotor cortices in each group. Figure 5 shows raster plots and Gaussian-convolved firing rates for exemplar channels in an exemplar rat from each group. In each group, exemplar channels show strong modulation of task-related multi-unit neural firing. Additionally, individual channels display distinct associations with different periods of the task including the approach/preparatory phase, reach-to-grasp execution, and post-grasp retrieval.

**Figure 5.**
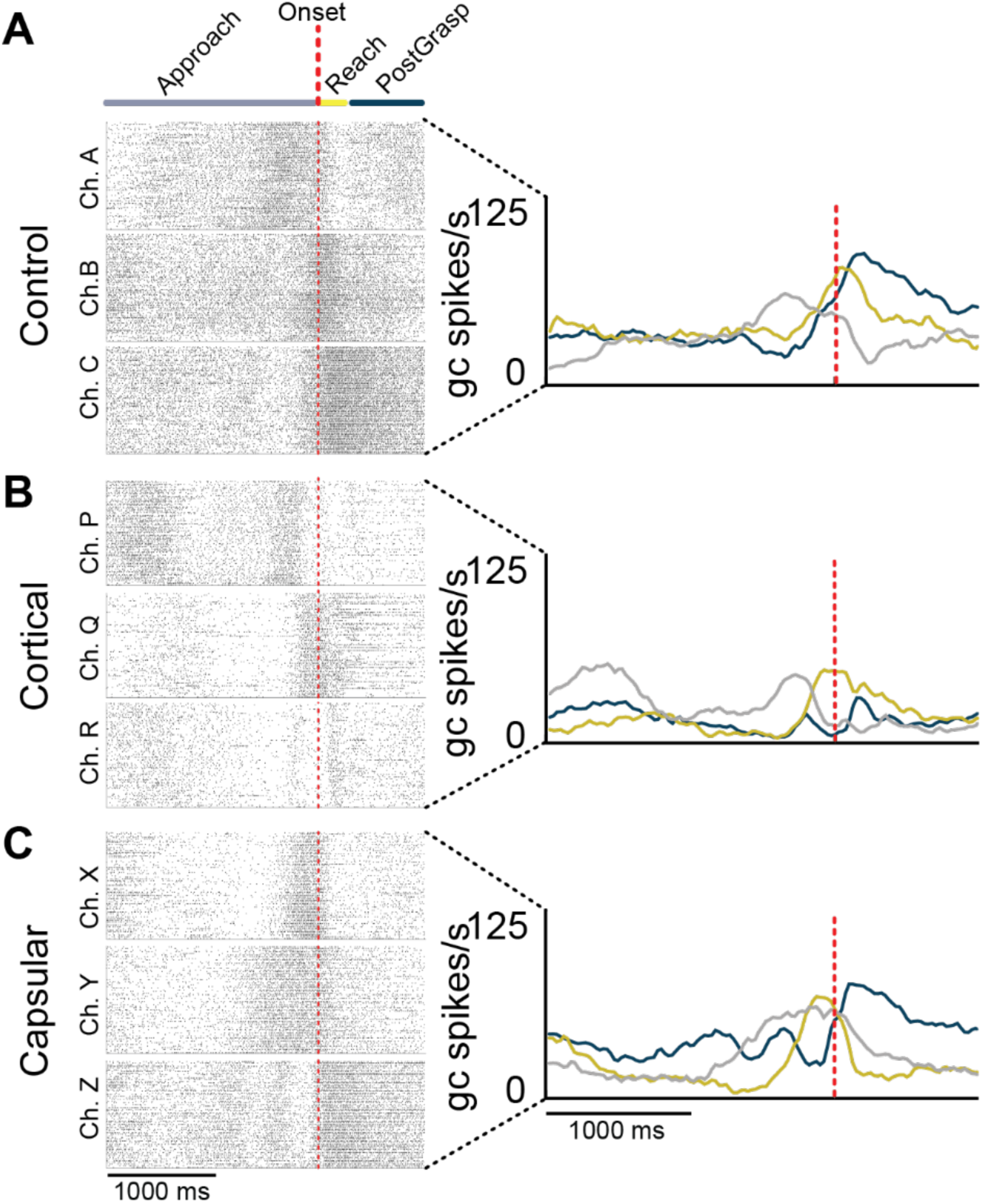
Task-related neural activity is present following post-infarct recovery. **A.** Reach-aligned spiking activity during task performance from three exemplar channels in a rodent within the control group. Increased multi-unit spiking can be seen across three different behavioral epochs including: Approach (planning), Reach execution, and PostGrasp (retrieval). Following smoothing with a Gaussian convolution, firing rates show the average activity across trials related to specific epochs. **B.** Spiking activity from three exemplar channels from a rodent in the cortical infarct group. **C.** Spiking activity from three exemplar channels from rodent in the capsular infarct group. Despite infarcts, following recovery task-related neural activity persists and can be observed in individual channels from non-infarcted sensorimotor regions in both stroke groups.

### 3.3 Lesion-specific alterations in global task-related firing

To examine the impact of each lesion model on the overall levels of task-related neural activity within each sensorimotor region after recovery, all multi-unit firing within each region was pooled across all channels within a region. Next, the mean Gaussian convolved firing rate on all successful trials across all days in all rodents within an infarct group was calculated. Figure 6 shows global task-related neural firing rate in each region and lesion model following recovery. In control animals, CFA exhibits strong task-related modulation centered around the onset of grasp. Additionally, RFA and S1 both exhibit task-related modulations that are lower in amplitude with distinct time courses. Specifically, activity in RFA is modulated over a wider interval spanning from reach onset through grasp onset, while S1 exhibits modulation of neural firing that peaks around grasp onset with maintained modulation extending into the post-grasp retrieval period. Following a focal infarct to CFA, task-related neural activity is disrupted in all three regions examined. Task-related neural activity is nearly abolished in CFA due to the lesion targeted directly to CFA (pairwise t-test Bonferroni and BH corrected: control v. cortical reach: p < 0.001, Cohen’s d =1.844; grasp: p < 0.001, Cohen’s d = 2.191 ; capsular v. cortical reach: p < 0.001, Cohen’s d = 1.262; grasp: p < 0.001, Cohen’s d = 1.532). In addition to decreases in task-related neural activity within the ischemic core and perilesional M1, cortical lesions also lead to a significant decrease in neural activity within RFA (pairwise t-test Bonferroni and BH corrected: control v. cortical reach: p < 0.001, Cohen’s d = 1.036; grasp: p < 0.001, Cohen’s d = 1.059 | capsular v. cortical reach: p < 0.001, Cohen’s d = 0.981; grasp: p < 0.001, Cohen’s d = 0.949) and to a lesser extent, S1 (pairwise t-test Bonferroni and BH corrected: control v. cortical reach: p < 0.001, Cohen’s d = 0.479; grasp: p < 0.001, Cohen’s d = 0.595 | capsular v. cortical reach: p < 0.001, Cohen’s d = 0.221; grasp: p < 0.001, Cohen’s d = 0.345). Finally, following recovery from a lesion to the internal capsule, task-related neural activity is also significantly decreased in each cortical region. While statistically significant changes are observed between neural firing in each lesion group and region due to the large number of channels pooled, the effect size is largest for the comparisons between the cortical lesion group and both the control and capsular lesion group are greater than the comparisons between the capsular lesion group and control group (pairwise t-test Bonferroni and BH corrected: RFA reach: p < 0.05, Cohen’s d = 0.136; grasp: p<0.001, Cohen’s d = 0.305 | CFA reach: p < 0.001, Cohen’s d = 0.821; grasp: p < 0.001, Cohen’s d = 0.414 | S1 reach: p < 0.001, Cohen’s d = 0.242; grasp p < 0.001, Cohen’s d = 0.300).

**Figure 6.**
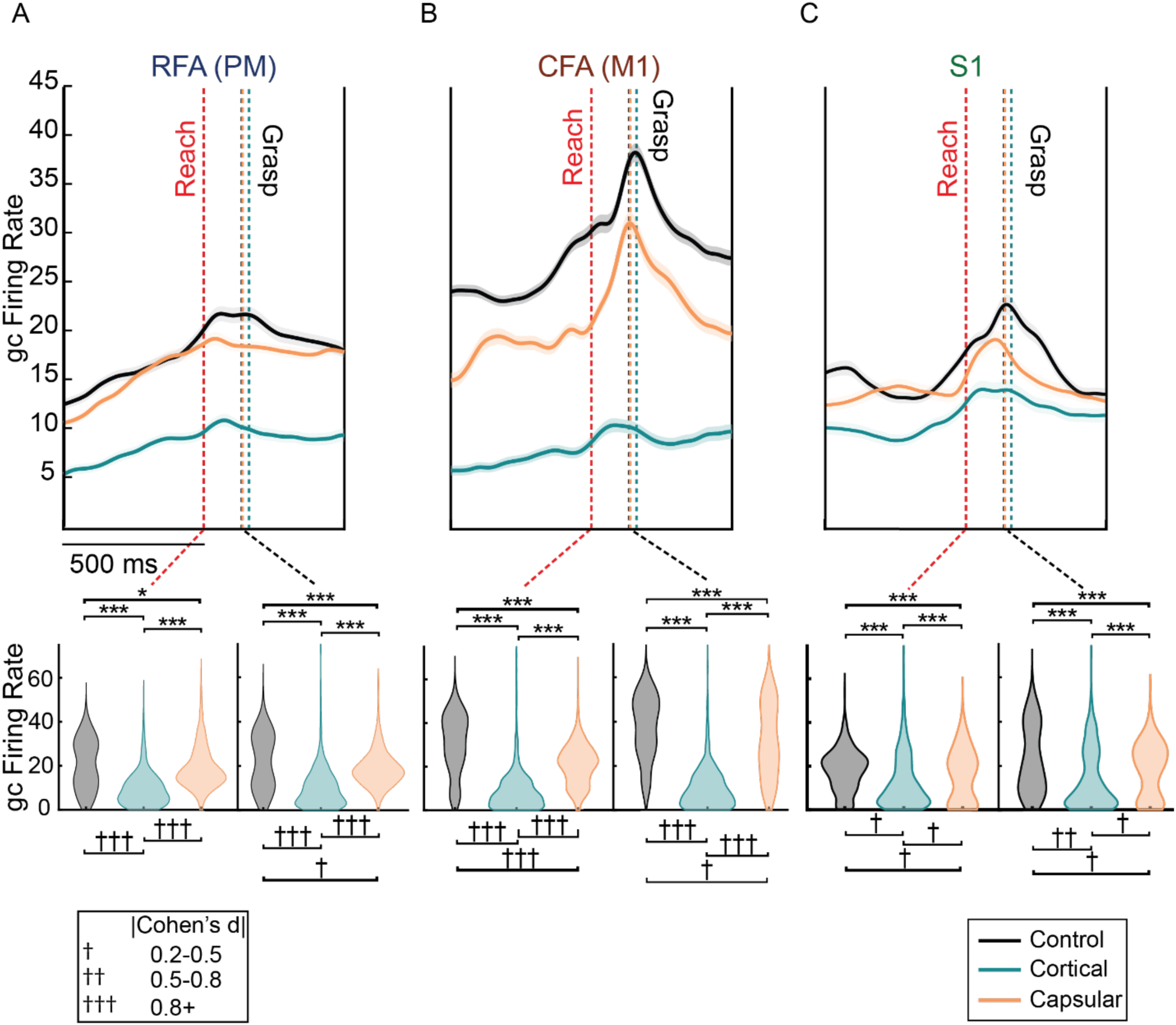
Region-specific alterations in task-related neural activity. **A.** Mean Gaussian convolved neural firing rate in RFA (PM) -500ms to +500ms from reach across all channels and rodents within each infarct group. Control rodents have greater neural firing than either experimental group. The degree of difference is smaller between control and capsular lesion groups (than when either group is compared to the cortical lesion group. **B.** Mean Gaussian convolved neural firing rates in CFA (M1) for each infarct group. The largest distribution difference between the control and capsular groups can be seen at reach, while the control and cortical groups are most different at grasp. Similarly, CFA activity in capsular and cortical infarct rats is most different at grasp with a larger effect size associated with the differences between capsular and cortical than cortical and control rats. **C.** Mean Gaussian convolved neural firing rates in S1 following each infarct. S1 shows the lowest overall difference between groups. Shaded regions indicate 95% CI across all trials in the plateau period for each rodent of a group. (* p < 0.05, ** p < 0.01, *** p < 0.001. † Cohen’s d = 0.2 - 0.5, †† Cohen’s d = 0.5-0.8, ††† Cohen’s d = 0.8+)

### 3.4 Lesion-specific alterations in task-related neural population activity

Along with changes in global neural activity, we also examined the recovery-related changes in the patterns of population-level neural activity across RFA, CFA, and S1. Figure 7 shows peri-event time histograms aligned to the onset of reach from exemplar rodents in each group. Firing rates were normalized (z-scored) relative to baseline firing rates. Channels were sorted by the time of peak firing rate and are color-coded by region for visualization. In control rats, task-related neural activity is distributed across regions. CFA activity is clustered around the reach and retrieval movements with activity in RFA and S1 occurring before, after, and temporally interspersed with neural activity in CFA. Following a cortical infarct to CFA, population-level neural activity is disrupted (Figure 7B). As expected, there are large disruptions in neural activity within and immediately around the lesion in CFA. Additionally, although task-related activations are observed in RFA and S1 following recovery, they are reduced in amplitude and spatial extent. In contrast, following recovery from an internal capsule infarct, despite some widening of the temporal window of task-related neural activity, the pattern of neural activity across the sampled ipsilesional neural population is maintained relative to healthy control animals (Figure 7C).

**Figure 7.**
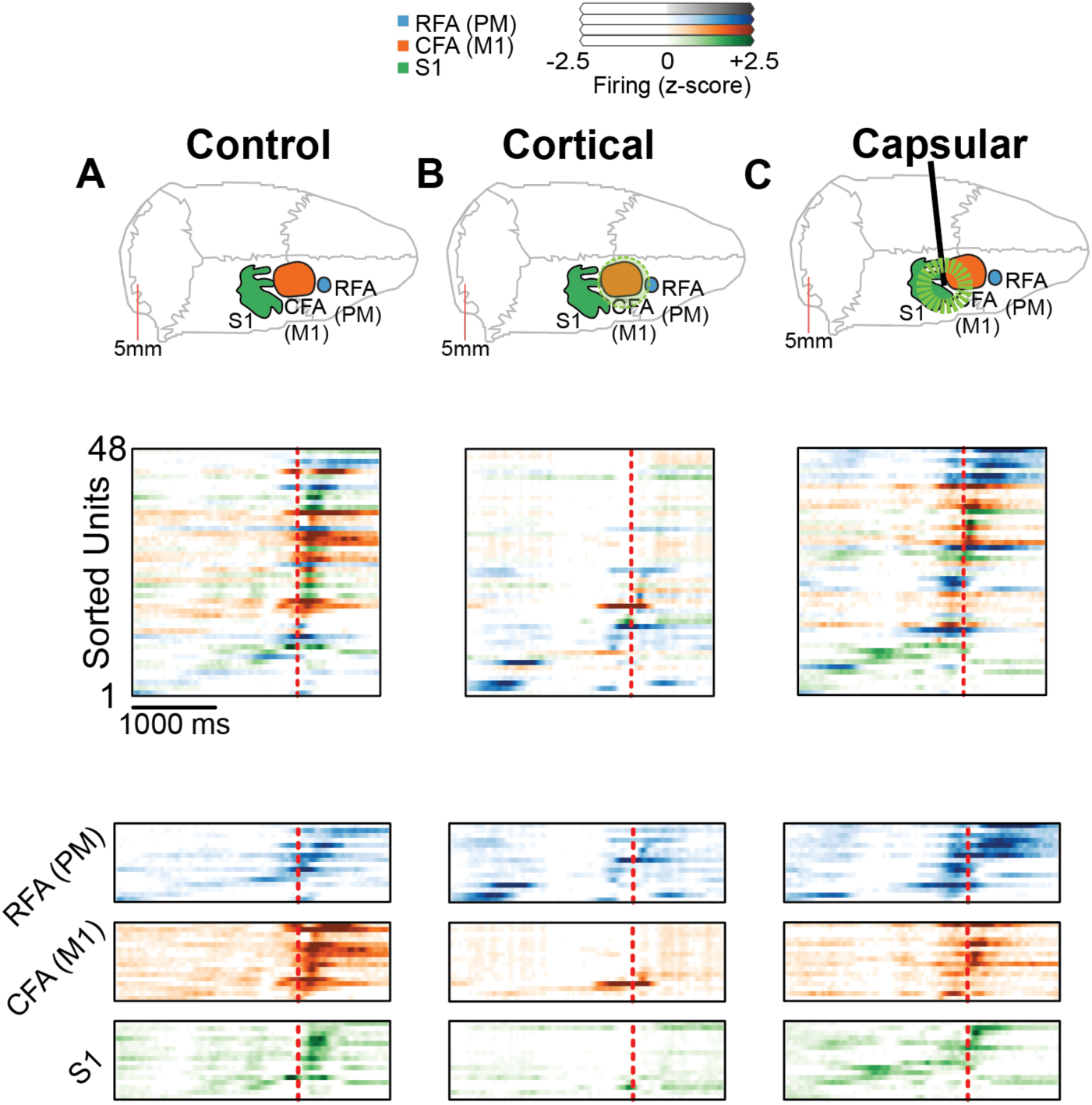
Exemplar movement-related population-level neural activity. Neural activity was aligned to the onset of reach, z-scored relative to baseline neural activity, and color-coded by region. Exemplar neural activity is shown for a single rat from each group with a schematic showing the lesion location (top) the entire population sorted by the time of peak neural activity (middle), and the neural activity subdivided by region (bottom) **A.** In a healthy control rodent consistent task-related neural activity is observed across regions with activations on individual channels in each region occurring interspersed throughout the reaching task. **B.** Following recovery from a cortical infarct to CFA, neural activity is nearly abolished in CFA. Task-related neural activity persists in RFA and S1 with decreased amplitude and spatial distribution of task-related neural activity. **C.** Following recovery from an internal capsule infarct, task-related neural activity is maintained with a similar interregional structure relative to healthy control rats.

### 3.5 Distinct prediction of task-related neural activity following cortical but not capsular infarcts

To examine the differences in the patterns of task-related population-level neural activity between regions and lesion groups, we trained a linear mixed model to predict neural activity on individual channels based upon differences in region, lesion model, and task epoch. To eliminate potential changes associated with the CFA array located within the lesion core or immediate perilesional region in the cortical lesion group, the model was trained using only channels in RFA and S1. The reference for model features was activity within the preparation/approach period in RFA in control rats. Therefore, positive and negative beta values indicate differences from this reference. The power of this method is that we can train the model using individual trials across animals and groups, increasing the power to detect differences in neural activity between lesion groups. Model coefficients are listed in the table and graphically displayed in Figure 8.

**Figure 8.**
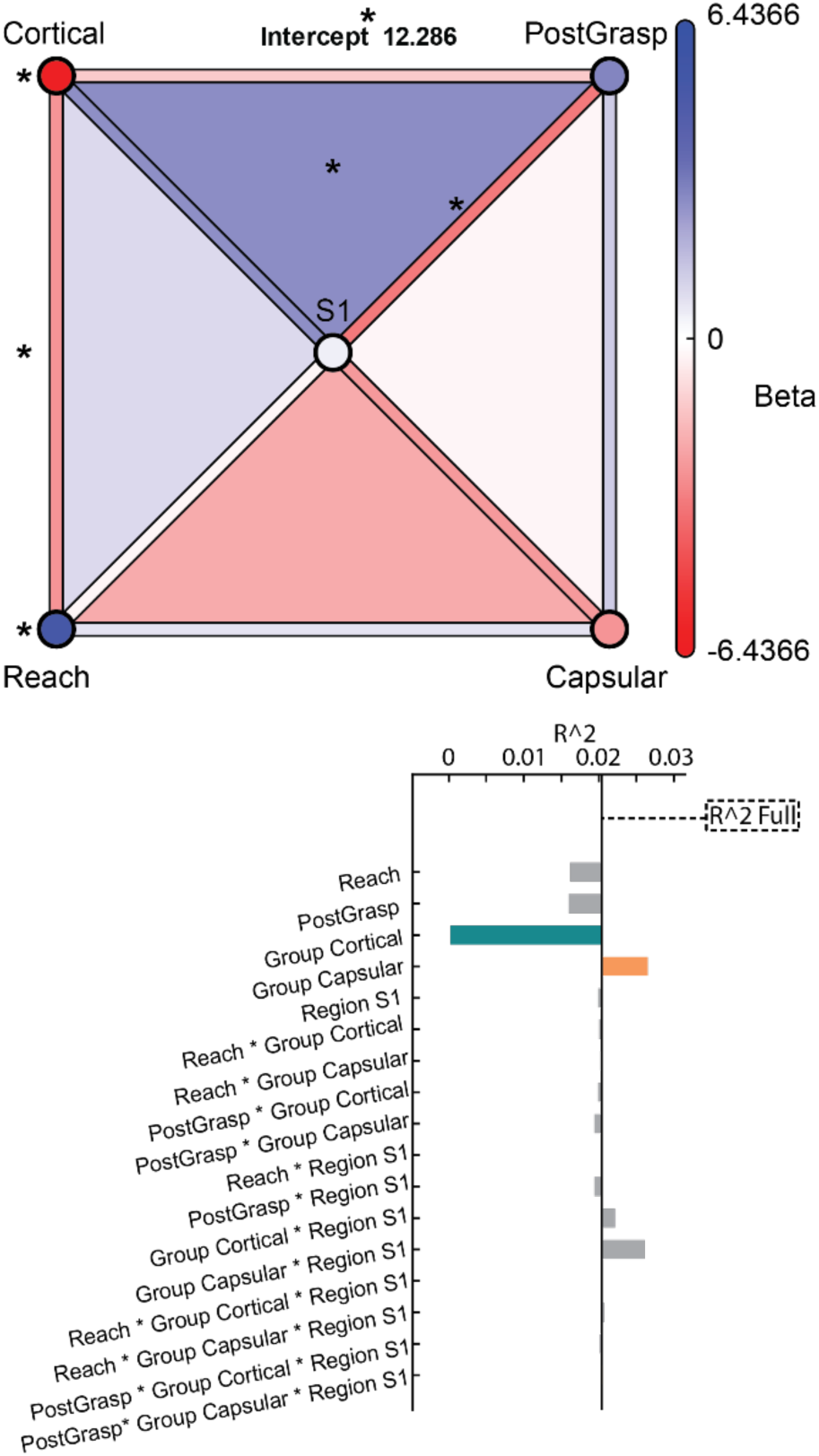
Generalized linear mixed model predicting neural activity. **A.** A Linear Mixed Model predicting neural firing across epoch, region, and lesion group. Data from CFA was excluded to avoid biases. Cortical injury rats have the lowest predicted neural firing compared to the reference (neural activity from control rats in RFA during the approach/preparation epoch). In contrast, there is no main effect for capsular lesion rats, indicating they do not have a significant modulation of their activity during the approach epoch (Intercept p < 0.001, β = 12.3; Cortical p = 0.044, β = -6.4; Capsular p = 0.40, β = -2.7). Each vertex represents a 1-dimensional main effect (epoch, region, or group), lines between vertices represent the 2-dimensional mixed effects, and faces of the pyramid represent the 3-dimensional mixed effects. **B.** To examine the impact of each effect on the model fit, the drop in neural variance explained by the model following a zeroing of the slope/Beta attributable to each fixed or mixed effect and retraining the model. With the removal of the cortical group effect and dependent mixed effects, the variance explained by the model drops, indicating a strongly reorganized pattern of neural activity in the cortical infarct group compared to the capsular infarct or control groups. (* p < 0.05)

**Table:** Generalized linear mixed model coefficients.

**Table: Generalized linear mixed model coefficients**
| Name | Estimate | SE | pValue |
| --- | --- | --- | --- |
| Intercept | 12.286 | 2.5212 | 1.10E-06 |
| Reach | 5.1962 | 0.8555 | 1.25E-09 |
| PostGrasp | 2.9834 | 0.8103 | 0.000232 |
| Cortical | -6.437 | 3.1896 | 0.043589 |
| Capsular | -2.715 | 3.1929 | 0.39519 |
| S1 | 0.5042 | 1.1573 | 0.6631 |
| Reach:Cortical | -2.684 | 1.083 | 0.013204 |
| Reach:Capsular | 0.7178 | 1.0869 | 0.50902 |
| PostGrasp:Cortical | -1.325 | 1.0258 | 0.19649 |
| PostGrasp:Capsular | 1.3558 | 1.0289 | 0.18759 |
| Reach:S1 | -0.287 | 1.2084 | 0.81223 |
| PostGrasp:S1 | -3.352 | 1.1439 | 0.003386 |
| Cortical:S1 | 2.7354 | 1.464 | 0.061688 |
| Capsular:S1 | -2.529 | 1.4644 | 0.084139 |
| Reach:Cortical:S1 | 1.2033 | 1.5294 | 0.43143 |
| Reach:Capsular:S1 | -2.64 | 1.5345 | 0.085359 |
| PostGrasp:Cortical:S1 | 3.4707 | 1.4476 | 0.016508 |
| PostGrasp:Capsular:S1 | -0.381 | 1.4514 | 0.79314 |

When examining the model output, rodents within the cortical group have the broadest impact in their task-related neural activity. There is a significant negative beta coefficient for the main effect associated with the cortical lesion group, indicating a decline in neural activity within the approach/preparation period relative to control animals (p = 0.044, β = -6.4). There is also a significant negative beta value for the mixed effect representing the cortical lesion group in the reach-to-grasp epoch, indicating an even greater decline in the task-related neural activity during the reach-to-grasp epoch for the cortical lesion group (p = 0.013, β = -2.7). In contrast, no fixed or mixed effects including the capsular group were statistically significant, indicating similar amplitudes of task-related neural activity in the control and capsular lesion group (Capsular main effect p = 0.40, β = -2.7). The only area where there is a significant positive modulation for the cortical group relative to the hierarchical structure is in S1 during the PostGrasp epoch (p = 0.017, β = 3.5). Therefore, the decline in firing in the cortical infarct group is reduced in S1 following grasp as compared to RFA or the approach/preparation and reach execution epochs. To examine the variance explained by each feature on the overall model performance, we then retrained the model with each main or interaction effect excluded. Removing the cortical group fixed effect causes a large relative decrease in the variance explained by the model (Figure 8). In contrast, removing the fixed effect associated with the capsular infarct group led to an increase in variance explained, suggesting that the lack of differences in neural dynamics between the control group and capsular infarct group leads to overfitting of the model (Figure 8).

## 4. Discussion

This study provides the first examination of the impact of lesion location on the specific patterns of task-related neural activity associated with cortical reorganization following stroke. Specifically, we compared patterns of ipsilesional task-related neural activity following focal infarcts targeting either the primary motor cortex or the internal capsule of rats. We initially hypothesized that each lesion group would show distinct patterns of neural reorganization relative to healthy controls. Although task-related neural activity was globally decreased in both lesion groups, surprisingly, the patterns of task-related neural activity across ipsilesional sensorimotor cortices were altered following lesions to the motor cortex but not following lesions to the internal capsule. While cortical reorganization has been examined extensively in preclinical stroke models, the most common preclinical models of stroke do not correspond to the lesion locations and extents typically seen in human patients. Therefore, the inclusion of an internal capsule lesion model allows us to specifically examine the clinical relevance of cortical reorganization to stroke recovery.

To our knowledge, this is the first time that direct recordings of task-related neural activity have been used to show that reorganization is dependent on the brain region most impacted by a focal infarct. Most striking is the lack of observed decline or reorganization following an infarct to the internal capsule. In particular, we used a linear mixed model to test the impact of lesion location on the prediction of neural firing. While the total variance explained by this approach can be modest due to the large number of factors impacting neural firing, the advantage of this approach is in harnessing the number of trials collected in a relatively small number of animals in order to have the power to test for the impact of group characteristics on neural firing. This modeling approach shows that there is a consistent structure in movement-related neural activity both in healthy animals and following recovery from a lesion to the internal capsule. This lack of reorganization is contrary to an extensive body of literature showing post-stroke reorganization in both preclinical models as well as clinical populations (Carter et al., 2010, 2012; Cramer et al., 1997; Fridman, 2004, 2004; Frost et al., 2003; Nelles et al., 1999; Nishibe et al., 2010; Nudo et al., 1996; Nudo & Milliken, 1996; Ward, 2003; Ward et al., 2003; Willer et al., 1993). Despite this evidence, when purely subcortical strokes are considered, recent studies have found a lack of evidence for recovery-related changes in resting state functional connectivity MRI (fcMRI) (Branscheidt et al., 2022; Makin & Krakauer, 2023). Our study builds on these examinations of subcortical stroke by providing a controlled examination of the impact of lesion location on neural firing in preclinical models with a specificity that is not possible in human patients. In particular, examining recovery following IC lesions is particularly relevant because the majority of patients with chronic motor deficits have some damage to subcortical structures with white matter damage particularly implicated in functional deficits (Corbetta et al., 2015; Edwardson et al., 2017; Murphy & Werring, 2020; Yang et al., 2022).

Although no ipsilesional reorganization was observed following internal capsular infarcts, cortical infarcts to CFA led to significant disruption of neural activity within RFA. Because RFA and CFA are strongly interconnected with reciprocal anatomic connections (Hira et al., 2013; Suter & Shepherd, 2015; Urban Iii et al., 2024; Z. Wang et al., 2018), this decrease in neural activity may represent diaschisis of the connected premotor area occurs following damage to primary motor areas. This decrease in neural activity following spontaneous recovery corresponds with decreased motor map area in RFA with spontaneous recovery following an infarct to CFA (Nishibe 2015). It is uncertain how rehabilitation would impact the decrease in task-related neural activity observed (Nishibe et al., 2010, 2015; Nudo & Milliken, 1996; Plautz et al., 2023).

Additionally, it is possible that cortical lesions extended beyond the CFA boundaries due to the overlapping and irregular patterns of blood vessels. Although we were not able to quantitatively test lesion size because of the implanted electrodes, lesions were targeted based upon stimulation maps of CFA and a collimated laser diode light source was used, allowing accurate targeting of focal cortical lesions to CFA. Cortical lesion locations and sizes were also confirmed by examining the locations of altered vasculature on the cortical surface using intraoperative photographs.

Along with differences in patterns of neural activity, the distinct recovery trajectories indicate the possible use of separate neural mechanisms driving recovery. Specifically, although both stroke groups have similar deficits immediately after lesioning and following recovery plateau, there was separation in the latency to the plateau in task success (∼28 days post-lesion) and ‘speed’ of movement between the two stroke groups, further suggesting that separate neural mechanisms drive recovery in each group. These results would align with the bimodal balance hypothesis which posits that motor system integrity determines the optimal mechanisms of functional recovery (Di Pino et al., 2014; Di Pino & Di Lazzaro, 2020). While the internal capsule infarct used does not directly damage motor cortex, as the primary output center of cortico-spinal projections (Basista & Yoshida, 2020; Dum & Strick, 1991; Dunkerley & Duncan, 1969; He et al., 1993, 1995; Lemon, 2008), this lesion would both damage corticospinal neurons originating in motor cortex through Wallerian degeneration as well as limit corticospinal outflow from all ipsilesional motor and premotor cortices. Therefore, recovery from an internal capsule lesion would be expected to require reorganization from potentially unobserved regions to enable descending motor signals to utilize alternative spinal projections. Additional studies will be important to test for the presence of reorganization in additional regions while varying both lesion location as well as corticospinal tract integrity.

Interestingly, despite the relationship between white matter damage and chronic motor deficits in clinical populations (Corbetta 2015), both or our infarct groups plateau around the same level of recovery. It is possible that recovery from capsular infarcts utilizes reorganization in either contralesional or ipsilesional regions outside of our areas of electrode coverage (Cai et al., 2016; Nelles et al., 1999). However, it is also possible that recovery from capsular infarcts in our rat model relies on extrapyramidal pathways that differ in strength between humans and rodents. Specifically, alternative avenues for recovery include extrapyramidal pathways such as the reticulospinal and rubrospinal tracts. In particular, the reticulospinal tract has broad projections and strong access to proximal/axial motor pools supporting rapid but less-fractionated movements and the rubrospinal/propriospinal circuits that may improve distal precision (Basista & Yoshida, 2020; Borrell et al., 2020; Brown, 1974; Fox, 1970; García-Alías et al., 2015; Liang et al., 2016; Mitchell et al., 2016). Additionally, the reticulospinal tract has been shown to be relevant for the production of movements following pyramidal tract damage in multiple species (Baker, 2011; Dewald et al., 1995; Fries et al., 1991; Zaaimi et al., 2012). Although conventional CST-indexed measures like ICMS maps and contralateral TMS-MEPs, which historically are used to show declines or alterations in motor maps, may underestimate descending drive within cortico-reticulospinal routes (Borrell et al., 2020; Cai et al., 2016; Nelles et al., 1999; Pineiro et al., 2001; Weiller et al., 1992; Willer et al., 1993), the recordings of task-related neural activity used here would be sensitive to both corticospinal and extrapyramidal corticofugal pathways. Additionally, even if extrapyramidal pathways facilitate recovery, the distinct patterns of reorganization between lesion groups highlights the importance of lesion-specific mechanisms of recovery.

Despite the important findings of lesion-specific patterns of post-stroke reorganization, there are several limitations to note. First, we were unable to perform quantitative evaluations of lesion volumes due to the presence of chronic electrode arrays in the immediate vicinity of cortical lesions. However potential inconsistencies in lesion size are limited due to the use of stimulation mapping for lesion targeting. Second, we examined the neural correlates of spontaneous recovery without modeling motor rehabilitation which is typically applied clinically. Furthermore, we also used young adult rodents (4-6 months old). Because most stroke survivors are older adults with multiple comorbidities, it is likely that we overestimated the level of spontaneous recovery expected (Martin et al., 2025). Additionally, rats were all pretrained on the task prior to lesions, therefore we were not testing the ability of rats to learn new tasks after stroke, and it is possible that overtraining could have impacted the progression of functional recovery. Finally, the analysis was focused on the time period after recovery plateaus to examine the differences in reorganization following recovery. It is possible that there may have been early reorganization in the internal capsule lesion group that occurred in the initial weeks after stroke; however, the limited number of trials limits the statistical power to examine the neural correlates of movement during the course of recovery.

Future analyses using Bayesian approaches may allow inferences into the progression of neural changes in the setting of a limited number of trials.

Our findings demonstrate the presence of distinct lesion-specific patterns of post-stroke neural activity and behavioral recovery trajectories. Moving forward, there are several important future directions to pursue. First, it will be critical to extend these findings into additional recording sites including the ipsilesional parietal cortex and contralesional sensorimotor cortices (Bundy et al., 2017; Cai et al., 2016; Pineiro et al., 2001).

Furthermore, examining the impact of rehabilitation on lesion-specific reorganization would provide additional insights into the relevance of cortical reorganization for functional recovery (Nudo et al., 1996; Nudo & Milliken, 1996.). Finally, it will be vital to test the lesion-specific impacts of targeted, open- and closed-loop neuromodulation therapies (Bundy et al., 2017; Guggenmos et al., 2013; Meyers et al., 2018), enabling the development of individualized, patient-specific approaches to stroke rehabilitation (Di Pino et al., 2014; Di Pino & Di Lazzaro, 2020; Murphy & Werring, 2020).

## Acknowledgements

Funding for this work was provided by the National Institutes of Health Grants R01NS030853, R01NS131466, F32NS100339, the Landon Center on Aging, and American Heart Association Grant 25CDA1455487.

## Data Availability

The data underlying the findings reported in this manuscript will be made available upon reasonable request to the corresponding author at the time of publication.

## References

Baker, S. N. (2011). The primate reticulospinal tract, hand function and functional recovery. The Journal of Physiology, 589(23), 5603–5612. 10.1113/jphysiol.2011.215160

Basista, M. J., & Yoshida, Y. (2020). Corticospinal Pathways and Interactions Underpinning Dexterous Forelimb Movement of the Rodent. Neuroscience, 450, 184–191. 10.1016/j.neuroscience.2020.05.050

Bauer, A. Q., Kraft, A. W., Wright, P. W., Snyder, A. Z., Lee, J.-M., & Culver, J. P. (2014). Optical imaging of disrupted functional connectivity following ischemic stroke in mice. NeuroImage, 99, 388–401. 10.1016/j.neuroimage.2014.05.051

Bergeron, M. (2003). Inducing photochemical cortical lesions in rat brain. *Current Protocols in Neuroscience*, Chapter 9, Unit 9.16. 10.1002/0471142301.ns0916s23

Blasi, F., Whalen, M. J., & Ayata, C. (2015). Lasting Pure-Motor Deficits after Focal Posterior Internal Capsule White-Matter Infarcts in Rats. Journal of Cerebral Blood Flow & Metabolism, 35(6), 977–984. 10.1038/jcbfm.2015.7

Borrell, J. A., Krizsan-Agbas, D., Nudo, R. J., & Frost, S. B. (2020). Effects of a contusive spinal cord injury on cortically-evoked spinal spiking activity in rats. Journal of Neural Engineering, 17(6), 066005. 10.1088/1741-2552/abc1b5

Branscheidt, M., Ejaz, N., Xu, J., Widmer, M., Harran, M. D., Cortés, J. C., Kitago, T., Celnik, P., Hernandez-Castillo, C., Diedrichsen, J., Luft, A., & Krakauer, J. W. (2022). No evidence for motor-recovery-related cortical connectivity changes after stroke using resting-state fMRI. Journal of Neurophysiology, 127(3), 637–650. 10.1152/jn.00148.2021

Brown, L. T. (1974). Rubrospinal projections in the rat. Journal of Comparative Neurology, 154(2), 169–187. 10.1002/cne.901540205

Bundy, D. T., Guggenmos, D. J., Murphy, M. D., & Nudo, R. J. (2019). Chronic stability of single-channel neurophysiological correlates of gross and fine reaching movements in the rat. PLOS ONE, 14(10), e0219034. 10.1371/journal.pone.0219034

Bundy, D. T., Souders, L., Baranyai, K., Leonard, L., Schalk, G., Coker, R., Moran, D. W., Huskey, T., & Leuthardt, E. C. (2017). Contralesional Brain–Computer Interface Control of a Powered Exoskeleton for Motor Recovery in Chronic Stroke Survivors. Stroke, 48(7), 1908–1915. 10.1161/STROKEAHA.116.016304

Cai, J., Ji, Q., Xin, R., Zhang, D., Na, X., Peng, R., & Li, K. (2016). Contralesional Cortical Structural Reorganization Contributes to Motor Recovery after Sub-Cortical Stroke: A Longitudinal Voxel-Based Morphometry Study. Frontiers in Human Neuroscience, 10. 10.3389/fnhum.2016.00393

Carter, A. R., Astafiev, S. V., Lang, C. E., Connor, L. T., Rengachary, J., Strube, M. J., Pope, D. L. W., Shulman, G. L., & Corbetta, M. (2010). Resting interhemispheric functional magnetic resonance imaging connectivity predicts performance after stroke. Annals of Neurology, 67(3), 365–375. 10.1002/ana.21905

Carter, A. R., Patel, K. R., Astafiev, S. V., Snyder, A. Z., Rengachary, J., Strube, M. J., Pope, A., Shimony, J. S., Lang, C. E., Shulman, G. L., & Corbetta, M. (2012). Upstream Dysfunction of Somatomotor Functional Connectivity After Corticospinal Damage in Stroke. Neurorehabilitation and Neural Repair, 26(1), 7–19. 10.1177/1545968311411054

Corbett, D., Carmichael, S. T., Murphy, T. H., Jones, T. A., Schwab, M. E., Jolkkonen, J., Clarkson, A. N., Dancause, N., Weiloch, T., Johansen-Berg, H., Nilsson, M., & Joy, M. T. (2017). Enhancing the Alignment of the Preclinical and Clinical Stroke Recovery Research Pipeline: Consensus-Based Core Recommendations From the Stroke Recovery and Rehabilitation Roundtable Translational Working Group*.

Corbetta, M., Ramsey, L., Callejas, A., Baldassarre, A., Hacker, C. D., Siegel, J. S., Astafiev, S. V., Rengachary, J., Zinn, K., Lang, C. E., Connor, L. T., Fucetola, R., Strube, M., Carter, A. R., & Shulman, G. L. (2015). Common Behavioral Clusters and Subcortical Anatomy in Stroke. Neuron, 85(5), 927–941. 10.1016/j.neuron.2015.02.027

Cramer, S. C., Nelles, G., Benson, R. R., Kaplan, J. D., Parker, R. A., Kwong, K. K., Kennedy, D. N., Finklestein, S. P., & Rosen, B. R. (1997). A Functional MRI Study of Subjects Recovered From Hemiparetic Stroke. Stroke, 28(12), 2518–2527. 10.1161/01.STR.28.12.2518

Dewald, J. P. A., Pope, P. S., Given, J. D., Buchanan, T. S., & Rymer, W. Z. (1995). Abnormal muscle coactivation patterns during isometric torque generation at the elbow and shoulder in hemiparetic subjects.

Di Pino, G., & Di Lazzaro, V. (2020). The balance recovery bimodal model in stroke patients between evidence and speculation: Do recent studies support it? Clinical Neurophysiology, 131(10), 2488–2490. 10.1016/j.clinph.2020.07.004

Di Pino, G., Pellegrino, G., Assenza, G., Capone, F., Ferreri, F., Formica, D., Ranieri, F., Tombini, M., Ziemann, U., Rothwell, J. C., & Di Lazzaro, V. (2014). Modulation of brain plasticity in stroke: A novel model for neurorehabilitation. Nature Reviews Neurology, 10(10), 597–608. 10.1038/nrneurol.2014.162

Dum, R., & Strick, P. (1991). The origin of corticospinal projections from the premotor areas in the frontal lobe. The Journal of Neuroscience, 11(3), 667–689. 10.1523/JNEUROSCI.11-03-00667.1991

Dunkerley, G. B., & Duncan, D. (1969). A light and electron microscopic study of the normal and the degenerating corticospinal tract in the rat. Journal of Comparative Neurology, 137(2), 155–183. 10.1002/cne.901370204

E. Mayo, Nancy, Wood-Dauphinee, Sharon, Ahmed, Sara, Carron, G., Higgins, Johanne, Mcewen, Sara, & Salbach, N. (1999). Disablement following stroke. Disability and Rehabilitation, 21(5–6), 258–268. 10.1080/096382899297684

Edwardson, M. A., Wang, X., Liu, B., Ding, L., Lane, C. J., Park, C., Nelsen, M. A., Jones, T. A., Wolf, S. L., Winstein, C. J., & Dromerick, A. W. (2017). Stroke Lesions in a Large Upper Limb Rehabilitation Trial Cohort Rarely Match Lesions in Common Preclinical Models. Neurorehabilitation and Neural Repair, 31(6), 509–520. 10.1177/1545968316688799

Eisner-Janowicz, I., Barbay, S., Hoover, E., Stowe, A. M., Frost, S. B., Plautz, E. J., & Nudo, R. J. (2008). Early and Late Changes in the Distal Forelimb Representation of the Supplementary Motor Area After Injury to Frontal Motor Areas in the Squirrel Monkey. Journal of Neurophysiology, 100(3), 1498–1512. 10.1152/jn.90447.2008

Favaretto, C., Allegra, M., Deco, G., Metcalf, N. V., Griffis, J. C., Shulman, G. L., Brovelli, A., & Corbetta, M. (2022). Subcortical-cortical dynamical states of the human brain and their breakdown in stroke. Nature Communications, 13(1), 5069. 10.1038/s41467-022-32304-1

Fox, J. E. (1970). RET1CULOSPINAL NEURONES IN THE RAT. Brain Research.

Fridman, E. A. (2004). Reorganization of the human ipsilesional premotor cortex after stroke. Brain, 127(4), 747–758. 10.1093/brain/awh082

Fries, W., Danek, A., & Witt, T. N. (1991). Motor responses after transcranial electrical stimulation of cerebral hemispheres with a degenerated pyramidal tract. Annals of Neurology, 29(6), 646–650. 10.1002/ana.410290612

Frost, S. B., Barbay, S., Friel, K. M., Plautz, E. J., & Nudo, R. J. (2003). Reorganization of Remote Cortical Regions After Ischemic Brain Injury: A Potential Substrate for Stroke Recovery. Journal of Neurophysiology, 89(6), 3205–3214. 10.1152/jn.01143.2002

Frost, S. B., Barbay, S., Mumert, M. L., Stowe, A. M., & Nudo, R. J. (2006). An animal model of capsular infarct: Endothelin-1 injections in the rat. Behavioural Brain Research, 169(2), 206–211. 10.1016/j.bbr.2006.01.014

Furuichi, Y., Maeda, M., Moriguchi, A., Sawamoto, T., Kawamura, A., Matsuoka, N., Mutoh, S., & Yanagihara, T. (2003). Tacrolimus, a Potential Neuroprotective Agent, Ameliorates Ischemic Brain Damage and Neurologic Deficits after Focal Cerebral Ischemia in Nonhuman Primates. Journal of Cerebral Blood Flow & Metabolism, 23(10), 1183–1194. 10.1097/01.WCB.0000088761.02615.EB

García-Alías, G., Truong, K., Shah, P. K., Roy, R. R., & Edgerton, V. R. (2015). Plasticity of subcortical pathways promote recovery of skilled hand function in rats after corticospinal and rubrospinal tract injuries. Experimental Neurology, 266, 112–119. 10.1016/j.expneurol.2015.01.009

Guggenmos, D. J., Azin, M., Barbay, S., Mahnken, J. D., Dunham, C., Mohseni, P., & Nudo, R. J. (2013). Restoration of function after brain damage using a neural prosthesis. Proceedings of the National Academy of Sciences, 110(52), 21177–21182. 10.1073/pnas.1316885110

Hakon, J., Quattromani, M. J., Sjölund, C., Tomasevic, G., Carey, L., Lee, J.-M., Ruscher, K., Wieloch, T., & Bauer, A. Q. (2018). Multisensory stimulation improves functional recovery and resting-state functional connectivity in the mouse brain after stroke. NeuroImage: Clinical, 17, 717–730. 10.1016/j.nicl.2017.11.022

He, S., Dum, R., & Strick, P. (1993). Topographic organization of corticospinal projections from the frontal lobe: Motor areas on the lateral surface of the hemisphere. The Journal of Neuroscience, 13(3), 952–980. 10.1523/JNEUROSCI.13-03-00952.1993

He, S., Dum, R., & Strick, P. (1995). Topographic organization of corticospinal projections from the frontal lobe: Motor areas on the medial surface of the hemisphere. The Journal of Neuroscience, 15(5), 3284–3306. 10.1523/JNEUROSCI.15-05-03284.1995

Hira, R., Ohkubo, F., Tanaka, Y. R., Masamizu, Y., Augustine, G. J., Kasai, H., & Matsuzaki, M. (2013). In vivo optogenetic tracing of functional corticocortical connections between motor forelimb areas. Frontiers in Neural Circuits, 7. 10.3389/fncir.2013.00055

Ikeda, S., Harada, K., Ohwatashi, A., Kamikawa, Y., Yoshida, A., & Kawahira, K. (2013). A New Non-Human Primate Model of Photochemically Induced Cerebral Infarction. PLOS ONE, 8(3), e60037. 10.1371/journal.pone.0060037

Jackson, G., & Chari, K. (2019). National Health Statistics Reports, Number 132, November 13, 2019. National Health Statistics Report.

Kang, D.-W., Chalela, J. A., Ezzeddine, M. A., & Warach, S. (2003). Association of Ischemic Lesion Patterns on Early Diffusion-Weighted Imaging With TOAST Stroke Subtypes. Archives of Neurology, 60(12), 1730. 10.1001/archneur.60.12.1730

Kim, H.-S., Hwang, J. H., Han, S.-C., Kang, G.-H., Park, J.-Y., & Kim, H.-I. (2021). Precision Capsular Infarct Modeling to Produce Hand Motor Deficits in Cynomolgus Macaques. Experimental Neurobiology, 30(5), 356–364. 10.5607/en21026

Kim, H.-S., Kim, D., Kim, R. G., Kim, J.-M., Chung, E., Neto, P. R., Lee, M.-C., & Kim, H.-I. (2014). A Rat Model of Photothrombotic Capsular Infarct with a Marked Motor Deficit: A Behavioral, Histologic, and microPET Study. Journal of Cerebral Blood Flow & Metabolism, 34(4), 683–689. 10.1038/jcbfm.2014.2

Kraft, A. W., Bauer, A. Q., Culver, J. P., & Lee, J.-M. (2018). Sensory deprivation after focal ischemia in mice accelerates brain remapping and improves functional recovery through Arc-dependent synaptic plasticity. Science Translational Medicine, 10(426), eaag1328. 10.1126/scitranslmed.aag1328

Lang, C. E., Waddell, K. J., Barth, J., Holleran, C. L., Strube, M. J., & Bland, M. D. (2021). Upper Limb Performance in Daily Life Approaches Plateau Around Three to Six Weeks Post-stroke. Neurorehabilitation and Neural Repair, 35(10), 903–914. 10.1177/15459683211041302

Lee, J.-K., Park, M.-S., Kim, Y.-S., Moon, K.-S., Joo, S.-P., Kim, T.-S., Kim, J.-H., & Kim, S.-H. (2007). Photochemically induced cerebral ischemia in a mouse model. Surgical Neurology, 67(6), 620–625. 10.1016/j.surneu.2006.08.077

Lemon, R. N. (2008). Descending Pathways in Motor Control. Annual Review of Neuroscience, 31(1), 195–218. 10.1146/annurev.neuro.31.060407.125547

Liang, H., Watson, C., & Paxinos, G. (2016). Terminations of reticulospinal fibers originating from the gigantocellular reticular formation in the mouse spinal cord. Brain Structure and Function, 221(3), 1623–1633. 10.1007/s00429-015-0993-z

Maeda, M., Moriguchi, A., Mihara, K., Aoki, T., Takamatsu, H., Matsuoka, N., Mutoh, S., & Goto, T. (2005). FK419, a Nonpeptide Platelet Glycoprotein IIb/IIIa Antagonist, Ameliorates Brain Infarction Associated with Thrombotic Focal Cerebral Ischemia in Monkeys: Comparison with Tissue Plasminogen Activator. Journal of Cerebral Blood Flow & Metabolism, 25(1), 108–118. 10.1038/sj.jcbfm.9600013

Maeda, M., Takamatsu, H., Furuichi, Y., Noda, A., Awaga, Y., Tatsumi, M., Yamamoto, M., Ichise, R., Nishimura, S., & Matsuoka, N. (2005). Characterization of a novel thrombotic middle cerebral artery occlusion model in monkeys that exhibits progressive hypoperfusion and robust cortical infarction. Journal of Neuroscience Methods, 146(1), 106–115. 10.1016/j.jneumeth.2005.01.019

Makin, T. R., & Krakauer, J. W. (2023). Against cortical reorganisation. eLife, 12, e84716. 10.7554/eLife.84716

Marin, M. A., & Carmichael, S. T. (2018). Stroke in CNS White Matter: Models and Mechanisms. Neuroscience Letters, 684, 193. 10.1016/j.neulet.2018.07.039

Martin, S. S., Aday, A. W., Allen, N. B., Almarzooq, Z. I., Anderson, C. A. M., Arora, P., Avery, C. L., Baker-Smith, C. M., Bansal, N., Beaton, A. Z., Commodore-Mensah, Y., Currie, M. E., Elkind, M. S. V., Fan, W., Generoso, G., Gibbs, B. B., Heard, D. G., Hiremath, S., Johansen, M. C., … on behalf of the American Heart Association Council on Epidemiology and Prevention Statistics Committee and Stroke Statistics Committee. (2025). 2025 Heart Disease and Stroke Statistics: A Report of US and Global Data From the American Heart Association. Circulation, 151(8). 10.1161/CIR.0000000000001303

Meyers, E. C., Solorzano, B. R., James, J., Ganzer, P. D., Lai, E. S., Rennaker, R. L., Kilgard, M. P., & Hays, S. A. (2018). Vagus Nerve Stimulation Enhances Stable Plasticity and Generalization of Stroke Recovery. Stroke, 49(3), 710–717. 10.1161/STROKEAHA.117.019202

Mitchell, E. J., McCallum, S., Dewar, D., & Maxwell, D. J. (2016). Corticospinal and Reticulospinal Contacts on Cervical Commissural and Long Descending Propriospinal Neurons in the Adult Rat Spinal Cord; Evidence for Powerful Reticulospinal Connections. PLOS ONE, 11(3), e0152094. 10.1371/journal.pone.0152094

Moon, S.-K., Alaverdashvili, M., Cross, A. R., & Whishaw, I. Q. (2009). Both compensation and recovery of skilled reaching following small photothrombotic stroke to motor cortex in the rat. Experimental Neurology, 218(1), 145–153. 10.1016/j.expneurol.2009.04.021

Murata, Y., & Higo, N. (2016). Development and Characterization of a Macaque Model of Focal Internal Capsular Infarcts. PLOS ONE, 11(5), e0154752. 10.1371/journal.pone.0154752

Murphy, S. JX., & Werring, D. J. (2020). Stroke: Causes and clinical features. Medicine, 48(9), 561–566. 10.1016/j.mpmed.2020.06.002

Nelles, G., Spiekermann, G., Jueptner, M., Leonhardt, G., Müller, S., Gerhard, H., & Diener, H. C. (1999). Reorganization of Sensory and Motor Systems in Hemiplegic Stroke Patients: A Positron Emission Tomography Study. Stroke, 30(8), 1510–1516. 10.1161/01.STR.30.8.1510

Nishibe, M., Barbay, S., Guggenmos, D., & Nudo, R. J. (2010). Reorganization of Motor Cortex after Controlled Cortical Impact in Rats and Implications for Functional Recovery. Journal of Neurotrauma, 27(12), 2221–2232. 10.1089/neu.2010.1456

Nishibe, M., Urban, E. T. R., Barbay, S., & Nudo, R. J. (2015). Rehabilitative Training Promotes Rapid Motor Recovery but Delayed Motor Map Reorganization in a Rat Cortical Ischemic Infarct Model. Neurorehabilitation and Neural Repair, 29(5), 472–482. 10.1177/1545968314543499

Nudo, R. J., & Milliken, G. W. (1996). Reorganization of movement representations in primary motor cortex following focal ischemic infarcts in adult squirrel monkeys.

Nudo, R. J., Wise, B. M., SiFuentes, F., & Milliken, G. W. (1996). Neural Substrates for the Effects of Rehabilitative Training on Motor Recovery After Ischemic Infarct. Science, 272(5269), 1791–1794. 10.1126/science.272.5269.1791

Paxinos, G., & Watson, C. (2013). The Rat Brain in Stereotaxic Coordinates (7th ed.). Elsevier.

Pineiro, R., Pendlebury, S., Johansen-Berg, H., & Matthews, P. M. (2001). Functional MRI Detects Posterior Shifts in Primary Sensorimotor Cortex Activation After Stroke: Evidence of Local Adaptive Reorganization? Stroke, 32(5), 1134–1139. 10.1161/01.STR.32.5.1134

Plautz, E. J., Barbay, S., Frost, S. B., Stowe, A. M., Dancause, N., Zoubina, E. V., Eisner-Janowicz, I., Guggenmos, D. J., & Nudo, R. J. (2023). Spared Premotor Areas Undergo Rapid Nonlinear Changes in Functional Organization Following a Focal Ischemic Infarct in Primary Motor Cortex of Squirrel Monkeys. The Journal of Neuroscience, 43(11), 2021–2032. 10.1523/jneurosci.1452-22.2023

Quattromani, M. J., Hakon, J., Rauch, U., Bauer, A. Q., & Wieloch, T. (2018). Changes in resting-state functional connectivity after stroke in a mouse brain lacking extracellular matrix components. Neurobiology of Disease, 112, 91–105. 10.1016/j.nbd.2018.01.011

Sawada, H., Nishimura, N., Suzuki, E., Zhuang, J., Hasegawa, K., Takamatsu, H., Honda, K., & Hasumi, K. (2014). SMTP-7, a novel small-molecule thrombolytic for ischemic stroke: A study in rodents and primates. Journal of Cerebral Blood Flow and Metabolism: Official Journal of the International Society of Cerebral Blood Flow and Metabolism, 34(2), 235–241. 10.1038/jcbfm.2013.191

Schrandt, C. J., Kazmi, S. S., Jones, T. A., & Dunn, A. K. (2015). Chronic Monitoring of Vascular Progression after Ischemic Stroke Using Multiexposure Speckle Imaging and Two-Photon Fluorescence Microscopy. Journal of Cerebral Blood Flow & Metabolism, 35(6), 933–942. 10.1038/jcbfm.2015.26

Song, H., Jung, W., Lee, E., Park, J.-Y., Kim, M. S., Lee, M.-C., & Kim, H.-I. (2017). Capsular stroke modeling based on somatotopic mapping of motor fibers. Journal of Cerebral Blood Flow & Metabolism, 37(8), 2928–2937. 10.1177/0271678X16679421

Sozmen, E. G., Hinman, J. D., & Carmichael, S. T. (2012). Models That Matter: White Matter Stroke Models. Neurotherapeutics, 9(2), 349. 10.1007/s13311-012-0106-0

Stolz, E., Jauss, M., Wessels, T., Wessels, C., Ellsiepen, A., & Reuter, I. (2006). In Determination of Stroke Etiology Contribution of Diffusion-Weighted Imaging.

Suter, B. A., & Shepherd, G. M. G. (2015). Reciprocal Interareal Connections to Corticospinal Neurons in Mouse M1 and S2. The Journal of Neuroscience, 35(7), 2959– 2974. 10.1523/JNEUROSCI.4287-14.2015

Tomizawa, A., Ohno, K., Jakubowski, J. A., Mizuno, M., & Sugidachi, A. (2015). Prasugrel reduces ischaemic infarct volume and ameliorates neurological deficits in a non-human primate model of middle cerebral artery thrombosis. Thrombosis Research, 136(6), 1224–1230. 10.1016/j.thromres.2015.09.013

Trautmann, E. M., Stavisky, S. D., Lahiri, S., Ames, K. C., Kaufman, M. T., O’Shea, D. J., Vyas, S., Sun, X., Ryu, S. I., Ganguli, S., & Shenoy, K. V. (2019). Accurate Estimation of Neural Population Dynamics without Spike Sorting. Neuron, 103(2), 292–308.e4. 10.1016/j.neuron.2019.05.003

Urban Iii, E. T., Hudson, H. M., Li, Y., Nishibe, M., Barbay, S., Guggenmos, D. J., & Nudo, R. J. (2024). Corticocortical connections of the rostral forelimb area in rats: A quantitative tract-tracing study. Cerebral Cortex, 34(2), bhad530. 10.1093/cercor/bhad530

Van Meer, M. P. A., Van Der Marel, K., Wang, K., Otte, W. M., El Bouazati, S., Roeling, T. A. P., Viergever, M. A., Berkelbach Van Der Sprenkel, J. W., & Dijkhuizen, R. M. (2010). Recovery of Sensorimotor Function after Experimental Stroke Correlates with Restoration of Resting-State Interhemispheric Functional Connectivity. The Journal of Neuroscience, 30(11), 3964–3972. 10.1523/JNEUROSCI.5709-09.2010

Wang, Y., Liu, G., Hong, D., Chen, F., Ji, X., & Cao, G. (2016). White matter injury in ischemic stroke. Progress in Neurobiology, 141, 45–60. 10.1016/j.pneurobio.2016.04.005

Wang, Z., Maunze, B., Wang, Y., Tsoulfas, P., & Blackmore, M. G. (2018). Global Connectivity and Function of Descending Spinal Input Revealed by 3D Microscopy and Retrograde Transduction. The Journal of Neuroscience, 38(49), 10566–10581. 10.1523/JNEUROSCI.1196-18.2018

Ward, N. S. (2003). Neural correlates of motor recovery after stroke: A longitudinal fMRI study. Brain, 126(11), 2476–2496. 10.1093/brain/awg245

Ward, N. S., Brown, M. M., Thompson, A. J., & Frackowiak, R. S. J. (2003). Neural correlates of outcome after stroke: A cross-sectional fMRI study. Brain, 126(6), 1430– 1448. 10.1093/brain/awg145

Watson, B. D., Dietrich, W. D., Busto, R., Wachtel, M. S., & Ginsberg, M. D. (1985). Induction of reproducible brain infarction by photochemically initiated thrombosis. Annals of Neurology, 17(5), 497–504. 10.1002/ana.410170513

Weiller, C., Chollet, F., Friston, K. J., Wise, R. J. S., & Frackowiak, R. S. J. (1992). Functional reorganization of the brain in recovery from striatocapsular infarction in man. Annals of Neurology, 31(5), 463–472. 10.1002/ana.410310502

Wen, T.-C., Sindhurakar, A., Ramirez, V. C., Park, H., Gupta, D., & Carmel, J. B. (2019a). Targeted Infarction of the Internal Capsule in the Rat Using Microstimulation Guidance. Stroke, 50(9), 2531–2538. 10.1161/STROKEAHA.119.025646

Wen, T.-C., Sindhurakar, A., Ramirez, V. C., Park, H., Gupta, D., & Carmel, J. B. (2019b). Targeted Infarction of the Internal Capsule in the Rat Using Microstimulation Guidance. Stroke, 50(9), 2531–2538. 10.1161/STROKEAHA.119.025646

Whishaw, I. Q., & Pellis, S. M. (1990). Reaching in the rat" a a single distal rotatory.

Willer, C., Ramsay, S. C., Wise, R. J. S., Friston, K. J., & Frackwiak, R. S. J. (1993). Individual patterns of functional reorganization in the human cerebral cortex after capsular infraction. Annals of Neurology, 33(2), 181–189. 10.1002/ana.410330208

Yamashita, Y., Wada, I., Horiba, M., & Sahashi, K. (2016). Influence of cerebral white matter lesions on the activities of daily living of older patients with mild stroke. Geriatrics & Gerontology International, 16(8), 942–947. 10.1111/ggi.12580

Yang, L., Li, M., Zhan, Y., Feng, X., Lu, Y., Li, M., Zhuang, Y., Lei, J., & Zhao, H. (2022). The Impact of Ischemic Stroke on Gray and White Matter Injury Correlated With Motor and Cognitive Impairments in Permanent MCAO Rats: A Multimodal MRI-Based Study. Frontiers in Neurology, 13, 834329. 10.3389/fneur.2022.834329

Zaaimi, B., Edgley, S. A., Soteropoulos, D. S., & Baker, S. N. (2012). Changes in descending motor pathway connectivity after corticospinal tract lesion in macaque monkey. Brain, 135(7), 2277–2289. 10.1093/brain/aws115

